# Open-pFind enables precise, comprehensive and rapid peptide identification in shotgun proteomics

**DOI:** 10.1101/285395

**Authors:** Hao Chi, Chao Liu, Hao Yang, Wen-Feng Zeng, Long Wu, Wen-Jing Zhou, Xiu-Nan Niu, Yue-He Ding, Yao Zhang, Rui-Min Wang, Zhao-Wei Wang, Zhen-Lin Chen, Rui-Xiang Sun, Tao Liu, Guang-Ming Tan, Meng-Qiu Dong, Ping Xu, Pei-Heng Zhang, Si-Min He

**Affiliations:** Key Laboratory of Intelligent Information Processing of the Chinese Academy of Sciences (CAS), Institute of Computing Technology, CAS, Beijing, China; University of Chinese Academy of Sciences, Beijing, China; National Institute of Biological Sciences, Beijing, Beijing, China; State Key Laboratory of Proteomics, Beijing Proteome Research Center, National Center for Protein Sciences (Beijing), Beijing Institute of Lifeomics, Beijing, China; State Key Laboratory of Biocontrol and Guangdong Provincial Key Laboratory of Plant Resources, College of Ecology and Evolution, Sun Yat-Sen University, Guangzhou, China

## Abstract

Shotgun proteomics has grown rapidly in recent decades, but a large fraction of tandem mass spectrometry (MS/MS) data in shotgun proteomics are not successfully identified. We have developed a novel database search algorithm, Open-pFind, to efficiently identify peptides even in an ultra-large search space which takes into account unexpected modifications, amino acid mutations, semi- or non-specific digestion and co-eluting peptides. Tested on two metabolically labeled MS/MS datasets, Open-pFind reported 50.5‒117.0% more peptide-spectrum matches (PSMs) than the seven other advanced algorithms. More importantly, the Open-pFind results were more credible judged by the verification experiments using stable isotopic labeling. Tested on four additional large-scale datasets, 70‒85% of the spectra were confidently identified, and high-quality spectra were nearly completely interpreted by Open-pFind. Further, Open-pFind was over 40 times faster than the other three open search algorithms and 2‒3 times faster than three restricted search algorithms. Re-analysis of an entire human proteome dataset consisting of ∼25 million spectra using Open-pFind identified a total of 14,064 proteins encoded by 12,723 genes by requiring at least two uniquely identified peptides. In this search results, Open-pFind also excelled in an independent test for false positives based on the presence or absence of olfactory receptors. Thus, a practical use of the open search strategy has been realized by Open-pFind for the truly global-scale proteomics experiments of today and in the future.

## INTRODUCTION

Shotgun proteomics has grown rapidly in recent decades, especially for peptide and protein identification^1^. Database search, which is based on searching tandem mass spectrometry (MS/MS) data against a proteome database, has long been the dominant approach^2^. However, more than 50% of MS/MS data acquired in shotgun proteomics have not been successfully identified^3^. For example, Chick *et al.* reported a large-scale dataset consisting of over one million spectra, in which only 35.4% were identified via SEQUEST^4^ at a 1% false discovery rate (FDR) for proteins^5^. Another dataset, recently proposed by Bekker-Jensen *et al.*, also contained over one million spectra, of which only 38.9% were identified via MaxQuant^6^, with a 1% FDR for proteins and peptides^7^.

As shown in a number of studies, restricted proteome search engines cannot identify peptides with unexpected modifications, which is a major reason underlying the low identification rate^8–11^. Therefore, a feasible solution involves enlarging the search space to retrieve more peptide candidates with any type of modification. Chick *et al.* demonstrated that database search using a large mass tolerance window of ±500 Da increased the identification rate from 35.4% to 45.5%, but it still failed to identify more than 50% of the spectra^5^. In addition, database search with such a large mass tolerance window is very time-consuming, and the time penalty for the increased search space was approximately 10‒100-fold^5, 12^. Recently, Kong *et al.* proposed a blind search algorithm, MSFragger^12^, that significantly improved the search speed compared with that of three other tested algorithms by using the ion index technique^13, 14^.

In addition to unexpected modifications, several other factors also hinder precise peptide identification, including semi- and non-specific digestion, in-source fragmentation and co-eluting peptides in mixed spectra^15, 16^. These factors are collectively called the *dark matter* of shotgun proteomics^12, 17^ and are uniformly treated as mass shifts^5, 12^. However, instead of mass shifts, the exact peptide forms, including reasonable types of modifications and enzymatic cleavage sites, should be determined, which will make the search space significantly larger. For example, peptides that were non-specifically digested *in silico* by trypsin were observed ∼160 times more frequently than common tryptic peptides, and modifications exponentially produce 100‒30,000 additional peptide candidates^18^. Furthermore, searching against such an ultra-large space may seriously hamper the accuracy of search engines because correct peptides are difficult to distinguish among vast numbers of random peptides. The target-decoy strategy and a few other approaches have been widely used for FDR estimation^19–21^; however, these methods are needed to independently verify the credibility of the results obtained via open search. These challenging problems have discouraged open search in routine use. Instead, restricted search engines are preferred in shotgun proteomics, although they usually yield a low identification rate.

In this study, we developed a novel algorithm, Open-pFind, which adopted a comprehensive and ultra-fast open search workflow. The search space of Open-pFind was significantly expanded, and key factors that affect the identification rate, including unexpected modifications, amino acid mutations, semi-/non-specific digestion and co-eluting peptides, were fully considered. The performance of Open-pFind was first evaluated with two metabolically labeled datasets. Open-pFind reported 50.5‒117.0% more peptide-spectrum matches (PSMs) than seven other database search engines, and more importantly, the results were more credible according to isotopic labeling experiments. With four other large-scale MS/MS datasets, the identification rate of Open-pFind was stable within a range of 70‒85% and was close to 100% for high-quality spectra. Finally, we re-analyzed an entire human proteome dataset consisting of ∼25 million spectra (hereafter referred to as the Kim data)^22^. More than one million peptides were identified, which were 86.7% more than those reported previously. A total of 12,723 genes were confidently identified within a 1% FDR at the protein level, and over 90% (11,536) were supported by at least three peptides, of which no olfactory receptors were found^23^. These results demonstrated that open search strategies, as made practical by Open-pFind, will most likely be the preferred tools for large-scale MS/MS data analyses in the future.

## RESULTS

### The workflow of Open-pFind

The workflow of Open-pFind consists of two steps: open search and restricted search (Fig. 1a). First, the MS data are preprocessed by pParse, in which multiple precursor ions corresponding to each tandem mass spectrum are calibrated and extracted^24^. Then, the MS/MS data are searched against the indexed database via the open search module (Fig. 1b). Next, a few key parameters, such as highly abundant modifications, enzymatic specificity and the mass deviation distribution of precursor ions, are automatically learned by the reranking procedure, which is similar to the widely used Percolator algorithm^19, 25^ but considers more features related to open search. Second, MS/MS data are searched again via the restricted module, which is similar to regularly used restricted engines, *e.g.*, SEQUEST^4^ and MaxQuant^6, 26^. However, the protein database is reduced, and highly abundant modifications are specified automatically; both of these processes are based on the information learned in the previous step rather than expert experience. The results obtained from both the open and restricted searches are merged and reranked again. Finally, PSMs, peptides and proteins are individually filtered according to the specified thresholds (*e.g.*, 1% FDR at each level).

**Fig. 1.**
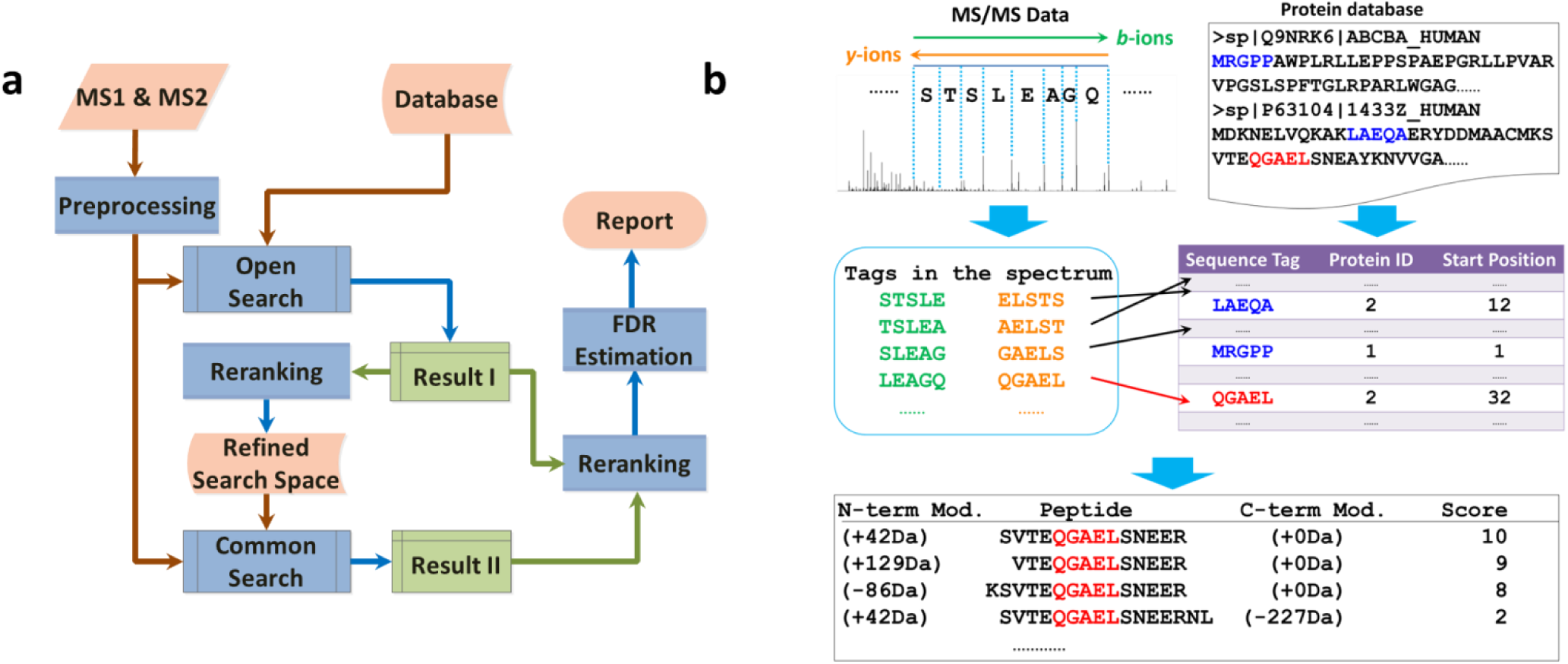
Workflow of Open-pFind, including the sub-workflow of the open search module. **a)** The workflow of Open-pFind. MS data are first preprocessed by pParse, and then the MS/MS data are searched by the open search module. Next, the MS data are re-searched by the restricted search module against the refined search space based on the learned information in the reranking step. Finally, the results obtained from both the open and restricted searches are merged, reranked again and reported. **b)** The default workflow of the open search module. For each spectrum, a few tags are extracted and then searched against the indexed protein database. Peptide candidates are then generated by extending the matched tags in proteins. Finally, peptides are scored with the spectrum and ranked; the mass shift between the precursor ion and each peptide is treated as a modification and localized by testing all valid positions.

The default sub-workflow of open search in Open-pFind is described as follows (Fig. 1b). A number of *k*-mer sequence tags are extracted from each spectrum and then retrieved in the indexed protein database. After finding the proteins matched with the tags, peptide candidates are generated by extending the matched tags in the proteins to fit the precursor ion mass, and a maximum of one non-zero mass shift was allowed when confirming the N- and C-termini of each peptide. The spectrum is then scored with each peptide, while the mass shift between the precursor ion and the peptide is treated as a modification. Open-pFind localizes all modifications in each peptide by testing all of the valid positions according to Unimod^27^, which is different from MSFragger that reports only peptides and unlocalized mass shifts. Finally, a number of high-score peptides are retained for each spectrum. A detailed description of the workflow is provided in the Online Methods.

### Open-pFind identified the highest number of PSMs, with half obtained from the extended search space

First, we evaluated the performance of Open-pFind with the metabolically labeled dataset Dong-Ecoli-QE (Fig. 2a and Supplementary Table 1). Open-pFind was compared with three open search engines, PEAKS^28, 29^, MODa^30^ and MSFragger^12^, as well as four restricted search engines, Comet^31^, Byonic^32^, MS-GF+^33^ and pFind^34, 35^ (Supplementary Tables 2 and 3).

**Fig. 2.**
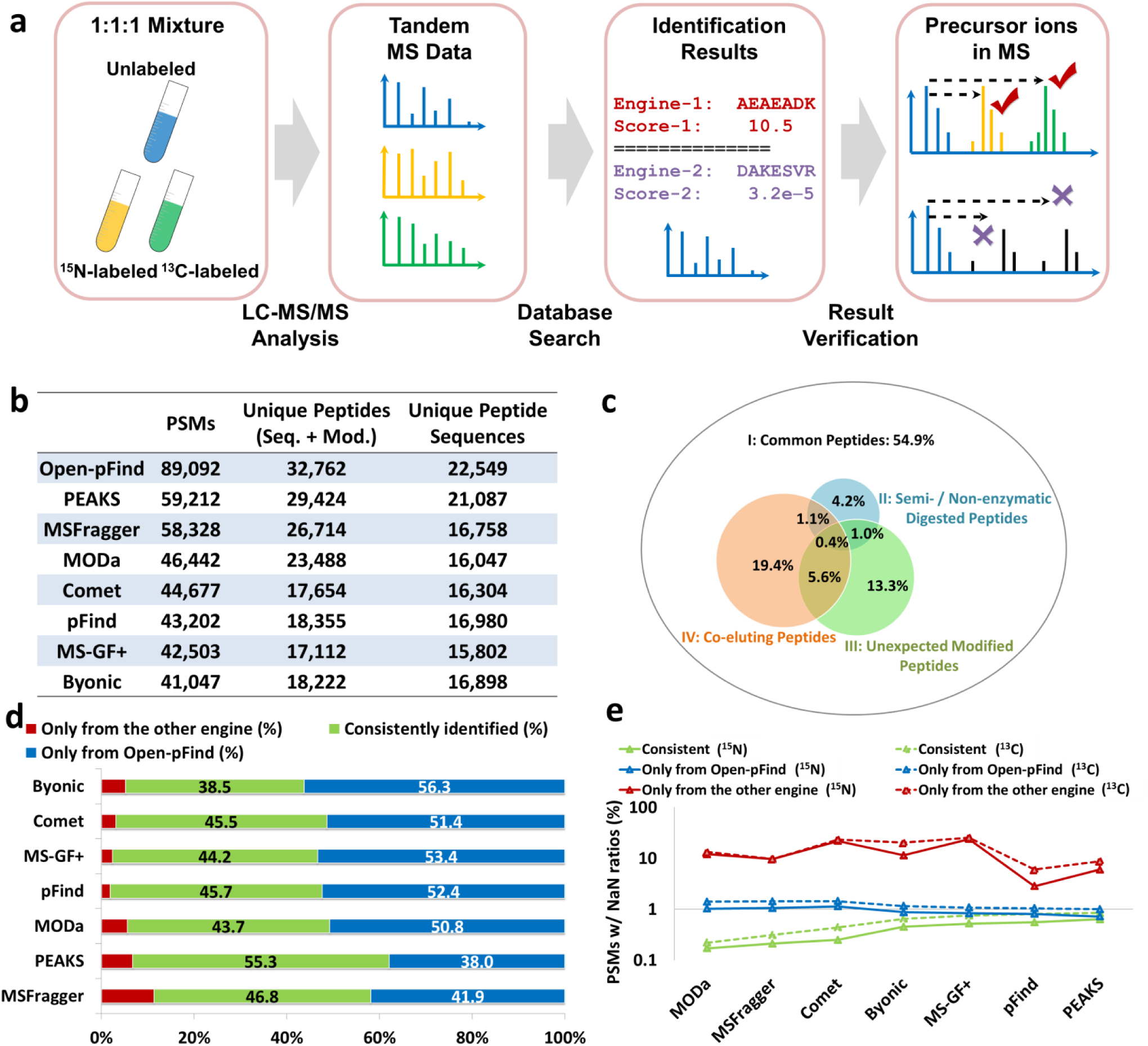
Performance evaluation of the Dong-Ecoli-QE dataset. **a)** Metabolically labeled datasets are searched against the protein database, in which only the non-labeled peptides are considered. Then, the accuracy is investigated by checking the percentage of the NaN-ratio PSMs. **b)** The numbers of identified PSMs, peptides and peptide sequences of each search engine. **c)** Distribution of the results of Open-pFind in different search spaces. **d)** The consistency of the results obtained by Open-pFind and each of the other search engines. **e)** The percentage of NaN-ratio PSMs in the consistently and separately identified results from Open-pFind and each of the other search engines.

Generally, the four open search engines reported more results than the restricted engines (Fig. 2b). Open-pFind identified 50.5% more PSMs, 11.3% more peptides (with modifications), and 6.9% more peptide sequences (regardless of modifications) than PEAKS, which ranked the second. A total of 54.9% of all PSMs identified via Open-pFind were obtained from the restricted search space, *i.e.*, corresponding peptides were also identified or at least surveyed by restricted search engines (Fig. 2c). In other words, 45.1% of the total PSMs were obtained from the extended search space produced by semi-/non-specific digestions, unexpected modification types, co-eluting peptides and the combination of these factors (Supplementary Note 1). When the PSMs identified by Open-pFind were combined with those of another search engine, Open-pFind identifications accounted for over 90% of the total PSMs in nearly every case, and 38‒56% of PSMs were uniquely reported by Open-pFind (Fig. 2d and Supplementary Table 4). Two PSMs were considered identical if both the peptide sequences (not distinguishing Leu and Ile) and modification types were the same.

### Metabolically labeled datasets were very helpful when evaluating search engine precision

Metabolically labeled datasets were designed and used to evaluate search engine precision in this study by using the percentage of PSMs with invalid quantitation values (Fig. 2a). Generally, if a spectrum is identified as an unlabeled peptide, the corresponding labeled precursor ions (^15^N- or ^13^C-labeled) should be observed in MS1 scans given the experimental design. In other words, if the labeled precursor ions of one peptide are not observed, which results in an invalid quantitation ratio (referred to as the *NaN ratio*, which is checked using pQuant^36^), the corresponding PSM is more likely to be incorrect. Consequently, the precision of the results obtained from different search engines using metabolically labeled datasets is evaluated by the percentage of PSMs or peptides associated with a NaN quantitation ratio.

The proportions of NaN-ratio PSMs (^15^N/^14^N or ^13^C/^12^C) in each part of the results shown in Fig. 2d were investigated. For the two blind search algorithms, MODa and MSFragger, only mass shifts rather than exact modification types were reported. Therefore, the exact numbers of N and C atoms in each peptide cannot be determined, making it impossible to locate the heavily labeled peptides in the MS1 spectra. As such, only the results with no modifications and those with common modifications (Supplementary Table 3) were used for the comparison shown in Fig. 2e.

Generally, among the PSMs identified by any two search engines, no more than 1% of them had a NaN ratio according to both metabolic labeling strategies, which was significantly less than the percentage of NaN-ratio PSMs in the uniquely identified results reported by one search engine. This finding also confirmed the widely accepted fact that PSMs identified consistently by different search engines are more credible (Fig. 2e). Among the PSMs uniquely identified by Open-pFind, the percentage of NaN-ratio PSMs were approximately 1%, only slightly higher than the percentage of NaN-Ratio PSMs in the overlapped results. In contrast, among the PSMs uniquely identified by any other search engine, the NaN-ratio PSMs exceeded 10% in most cases. A similar but wider comparison between any two engines indicated that open search engines reported more precise results than the restricted engines in terms of the peptides in the restricted search space (Supplementary Fig. 1), a finding that was also confirmed by Kong *et al*^12^. However, if all PSMs with any types of modifications were considered, the percentages of NaN-ratio PSMs identified via open search engines sharply increased, especially for the two blind search engines (Supplementary Fig. 2). In this context, it is worth noting that the Open-pFind results had the second smallest percentage of NaN-ratio PSMs, second only to pFind, indicating that the PSMs from the extended search space were also highly precise. Two additional analyses confirmed the above conclusions: one analysis utilized another metabolically labeled dataset, Xu-Yeast-QEHF (Supplementary Fig. 3), and the other analysis utilized the same dataset, Dong-Ecoli-QE, with three engines: Open-pFind, MaxQuant and SEQUEST-HT (Supplementary Note 2).

The identified proteins were directly inferred from the identified peptides; each peptide was uniquely matched to one protein in the database. For the proteins consistently identified by eight engines, Open-pFind reported 12.8‒94.3% more peptides on average (Supplementary Table 5), which is an important characteristic denoting stability and reproducibility in quantitative proteomics experiments. The percentage of NaN-ratio proteins was also an important indicator of the precision of protein identification. Similar to analysis at the PSM level, very few proteins with NaN ratios were found in the consistent results (Supplementary Fig. 4). Comparing the proteins uniquely identified by Open-pFind and those consistently identified by Open-pFind and another search engine, the percentages of NaN proteins are similar, especially for proteins with at least two supporting peptides. In contrast, among the proteins uniquely identified by any of the other search engines, the proportions of NaN-ratio proteins were 3‒30 times larger than those found in the Open-pFind results.

### Learning from the metabolic labeling technique

The metabolic labeling technique is also helpful in revealing why spectra are misidentified via different search engines and for improving search engine precision. Generally, a spectrum with a NaN-ratio peptide reported by one search engine may be identified as a different normal-ratio peptide by another search engine. As described above, the normal-ratio peptide is more likely to be a correct identification. Thus, for the former search engine, this could be used to optimize the scoring function. For all NaN-ratio PSMs from Open-pFind, only less than 10% were revived by other engines, *i.e.*, identified as normal-ratio peptides (Supplementary Fig. 5). In contrast, Open-pFind revived ∼40% of NaN-ratio PSMs reported by other search engines.

For the open search engines, Open-pFind reported an overlapping peptide different from the one reported by the other engine for ∼90% of the revived spectra (Supplementary Table 6): specifically, a peptide identified via Open-pFind appeared in that of the other engine or vice versa (*e.g.*, TAEHVAK/EHVAK is a pair of overlapping peptides). In other words, these results from the other open search engines were *partially* correct, while Open-pFind confirmed the exact termini of the peptides and modification types, as well as the precise precursor information. For example, Open-pFind reported a C-terminal-specific peptide carbamyl-GAAGGIGQALALLLK with an N-terminal carbamylation (*P*_1_) for the spectrum shown in Supplementary Fig. 6a, while MSFragger reported an overlapping tryptic peptide ***VAVL***GAAGGLGQALALLLK with a mass shift of –337.3114 Da (*P*_2_). However, the actual mass difference of these two peptides (*P*_2_ – *P*_1_) was 339.2522 Da. This result implied that the mass shift of –337.3114 Da reported by MSFragger did not represent a real modification because a ∼2-Da mass difference existed between the initially exported precursor ion and the actual one confirmed by Open-pFind (Supplementary Fig. 6b). This finding also demonstrated that exact precursor ions were very important for the confirmation of modification types.

In terms of the restricted search engines, over 90% of peptides reported by MS-GF+ and Comet were *partially* correct, which was similar to the behavior of the open search engines (Supplementary Table 6). However, this number was lower for Byonic and pFind. Byonic adopted a different protein FDR control strategy that a few low-quality PSMs from reliable proteins might be reported (Online Methods). An example in Supplementary Figs. 6c-f shows the differences between Open-pFind and the restricted search engines. For the same spectrum, Open-pFind reported a tryptic peptide with a deamidation, while MS-GF+ and Comet reported the unmodified form of this peptide, which obviously matched fewer fragment ions. Byonic reported a completely different peptide, which matched few peaks in the spectrum. The isotopic envelopes of the unlabeled peptide reported by Open-pFind, as well as the corresponding ^15^N- and ^13^C-labeled forms shown in MS1, matched the theoretical values precisely. In contrast, the monoisotopic precursor ions of the other two identifications had larger mass deviations, which resulted in invalid quantitation values (Supplementary Fig. 6f). This example indicated again that peptides reported by Open-pFind were more accurate, and more importantly, the metabolic labeling technique is extremely helpful when distinguishing correct individual PSMs, which will facilitate the improved design of search engines.

### Open-pFind yielded a high and stable identification rate with different types of large-scale datasets

Four large-scale, previously published datasets were used in this section: Mann-Human-Velos^37^, Gygi-Human-QE^5^, Mann-Mouse-QEHF^38^ and Pandey-Human-Elite^22^ (Supplementary Table 1). For all of the datasets, Open-pFind yielded the highest identification rate (77% on average), while for the seven other search engines the identification rates varied from 50‒65% (Fig. 3a). The numbers of target and decoy peptides obtained by Open-pFind with lower scores were almost identical (Supplementary Fig. 7), which proved that the algorithm had no bias between the target and decoy databases^39^. Similar to the analysis of the Dong-Ecoli-QE dataset, 30‒45% of the identifications involved in the extended search space were not identified via restricted search engines (data not shown).

**Fig. 3.**
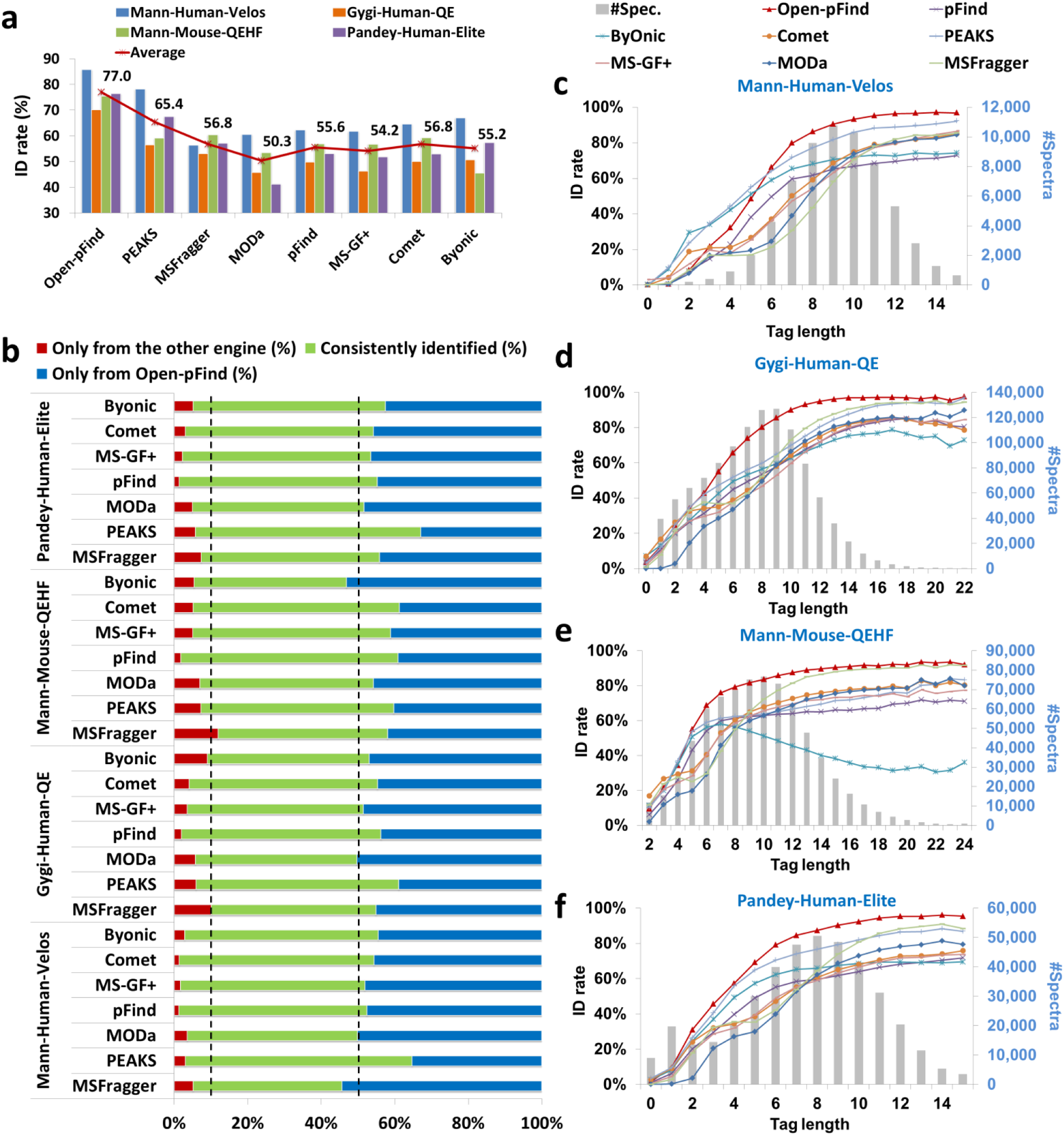
Performance evaluations of the four published datasets. **a)** The identification rate of each engine with the four datasets. **b)** The consistency of the results obtained by Open-pFind and each of the other search engines. The two dotted lines denote 10% and 50% on the x-axis. **c-f)** Analyses of the unidentified spectra in the four datasets. The curves denote the identification rates of the spectra with different tag lengths, and the histograms denote the distribution of the number of the total spectra at each tag length.

For all four datasets, Open-pFind almost always covered over 90% of any union sets of PSMs identified via itself and each of the other search engines (Fig. 3b and Supplementary Table 4). However, for any two restricted engines, the corresponding percentages varied from 60‒90%. Tessier *et al*. also confirmed this conclusion and proposed that although a few key parameters were identical for restricted engines, the *real* search spaces were still quite different, leading to disagreements among search engines^40^. Therefore, a complete or consistent search space is essential in yielding a stable identification rate, in addition to identifying more PSMs and peptides. We also compared the results of Open-pFind with those reported in previous studies. For example, with the Gygi-Human-QE dataset, Open-pFind covered 92.2% of the results reported by Chick *et al*. and reported 32.1% more PSMs with the same FDR threshold of ∼0.1% at the PSM level.

Search engine precision for these four datasets was also evaluated in this study. Although these datasets were not metabolically labeled, and thus the quantitative values could not be used to evaluate search engine precision, the entrapment strategy^41^ can be applied as an alternative approach. The decrease in the identification rate of pFind was 2‒3 times higher than that of Open-pFind, suggesting that the designed scheme of Open-pFind was more stable (Supplementary Note 3). In addition, although the same FDR threshold was controlled, more correct peptides from the authentic database, rather than the entrapment strategy, were obtained by Open-pFind.

### Nearly 100% of high-quality spectra are identified with a complete search space

We also investigated why a few spectra remained uninterpretable for Open-pFind. First, spectra are classified according to the lengths of their longest tags, which are treated as a feature related to spectral quality. For example, a 0-length tag indicates that no mass difference from any two peaks is equal to one amino acid residue within a given fragment ion tolerance. A spectrum with a longer tag meant that it was more likely to have been formed by a real peptide because more fragmentation information was provided. Generally, the identification rates of spectra with longer tags were higher for all engines (Figs. 3c-f). For all four datasets, the identification rate of Open-pFind was always greater than 90% and even close to 100% for spectra with tags longer than ten, suggesting that the search space of Open-pFind is close to complete for routine MS/MS data analysis. Additionally, the scoring scheme of Open-pFind effectively distinguishes correct peptides from the random peptides, even in such an ultra-large search space.

The identification rates of Byonic sharply decreased when spectra with longer tags were considered in the Mann-Mouse-QEHF dataset (Fig. 3e), likely because more large-mass peptides were present in this dataset, and their precursor ions were not correctly exported. Among all PSMs identified via Open-pFind in this dataset, 55.0% of their precursor ions were larger than 1,500 Da, of which only 50.1% were correctly exported by the vendor’s software. However, in the other datasets, the proportion of precursor ions larger than 1,500 Da was markedly less, for example, only 38.8% for the Pandey-Human-Elite dataset, and 82.1% of which were extracted correctly by the vendor’s software. Spectra with incorrectly assigned precursor ions cannot be matched to correct peptides. We also tested pFind using the precursor ions extracted by vendor software rather than pParse, and the distribution of identification rates was similar to that of Byonic (Supplementary Fig. 8), which again proved that extracting correct precursor ion masses was very important for search engine design.

### The speed of Open-pFind was comparable to or even faster than that of restricted engines with a ∼10^5^ times larger search space

The running times of the eight search engines for all six datasets in Supplementary Table 1 were comprehensively analyzed. Compared with the three other open search engines, Open-pFind was on average more than 40 times faster than MSFragger, MODa and PEAKS (Fig. 4a and Supplementary Table 7). Restricted search engines were generally faster than the open search engines, which was reasonable given the much smaller search space. As shown in Fig. 4b, the search space of Open-pFind was five orders of magnitude larger than the space considering only fully specific digestion with common modifications. However, Open-pFind was only slightly slower than pFind and 2‒3 times as fast as the three other restricted search engines (Byonic, Comet and MS-GF+), owing to the efficient tag-based workflow and the reduction of the database after open search (Online Methods). For the Xu-Yeast-QEHF dataset, Open-pFind was approximately one time faster than pFind because a six-frame-translated database was used, in which most proteins were irrelevant to all spectra. Therefore, removing these proteins according to the open search results was beneficial for speeding up the subsequent restricted search.

**Fig. 4.**
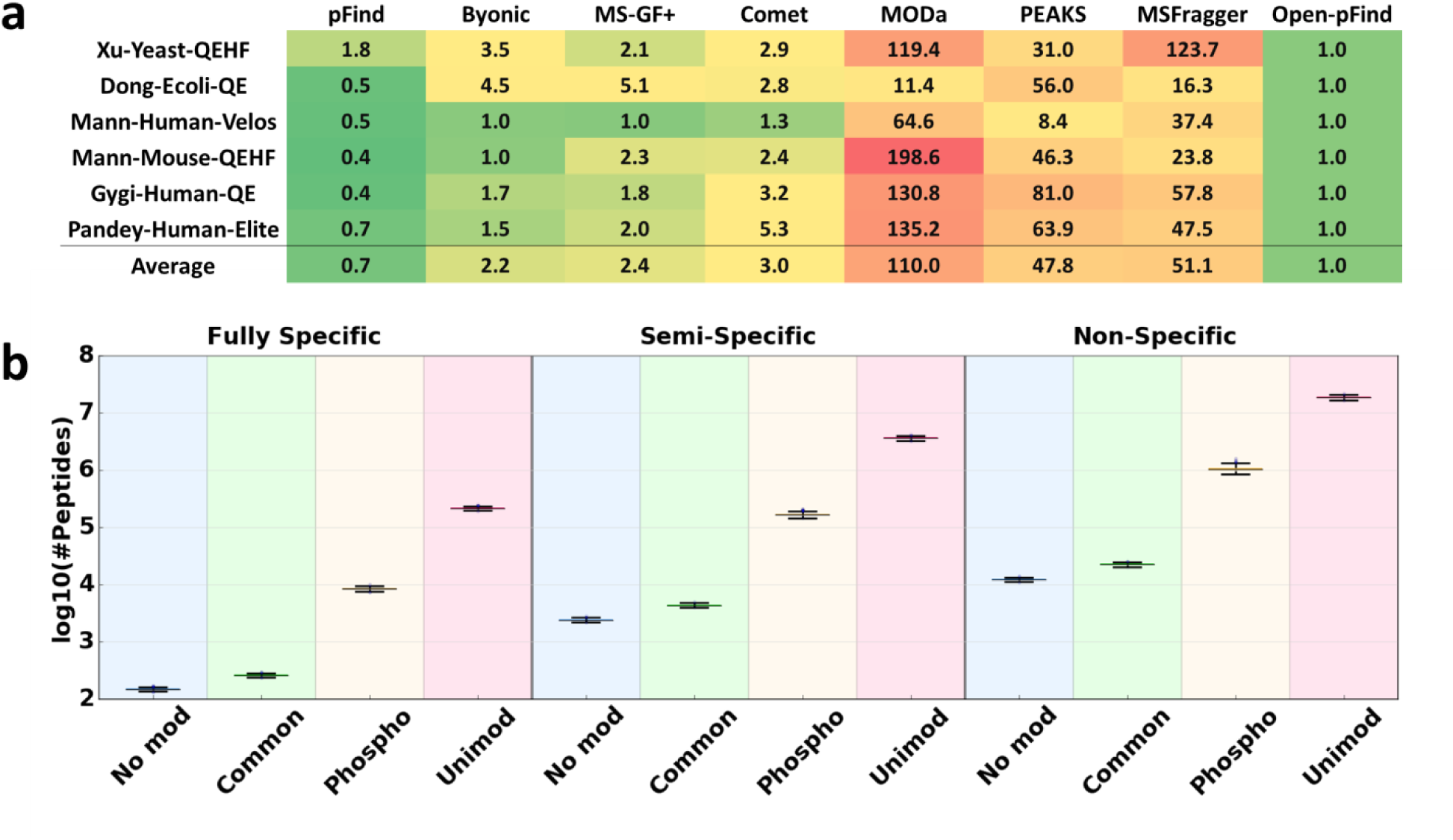
Comparison of the speed of each engine for various datasets. **a)** The normalized running times of the eight search engines. **b)** Boxplots showing the search spaces for different search modes, including a search with no modifications (No mod), restricted search (Common, with carbamidomethylation of C, oxidation of M, Gln→pyro-Glu at N-termini of peptides and acetylation at N-termini of proteins), phosphorylation search (Phospho, with common modifications and phosphorylation of S, T, and Y) and open search (Unimod, with at most one modification in Unimod), together with three different types of digestion. For each mode, 1,000 experiments were performed, each with 1,000 randomly selected proteins from UniProt database, which were digested *in silico* into peptides.

We further benchmarked the running times of the four open search engines using one raw data file of 41,820 spectra from Gygi-Human-QE^5, 12^ using the smaller, reviewed UniProt^42, 43^ human protein database (approximately one-eighth of the reviewed and unreviewed database of proteins) (Supplementary Table 8). Open-pFind was 14.4 and 16.0 times faster than PEAKS and MODa, respectively, and twice as fast as MSFragger when the search was restricted to fully tryptic peptides. When the search was extended to semi- and non-tryptic peptides were considered, the running time of Open-pFind nearly remained the same, whereas those of the other three search engines increased to varying degrees. We also tested Open-pFind in the open search mode with two unexpected modifications and in the blind search as in MODa and MSFragger (Online Methods); the Open-pFind search times were only 1.0‒3.4 times greater than those of the default workflow, and the identification rate slightly decreased because more irrelevant peptides were randomly matched to the spectra. Another analysis of the *T. tengcongensis* dataset^14, 44^ (Supplementary Table 9) confirmed the same conclusion: Open-pFind was at least ten times as fast as the other algorithms.

### Comprehensive analysis of the Kim data

It took Open-pFind 3,674 minutes (∼60 h) on a common PC, or 282 min (∼5 h) on a 64-core 64 GB RAM workstation, to search all of the ∼25 million spectra in the Kim data^22^. The target-decoy database used in this study contained 305,558 of both reviewed and unreviewed protein sequences (111.371 MB in total). In other words, Open-pFind processed ∼113 spectra per second, suggesting that peptide and protein identification is not a bottleneck, even with such an ultra-large-scale dataset using a common PC. The average identification rate was 62.5% for all 85 samples, and over 70% spectra were identified for the in-gel digested samples analyzed on an LTQ Orbitrap Velos MS (Fig. 5a). In addition, all peptides identified in the 85 samples were further filtered with a 1% FDR threshold at the peptide level. A total of 548,371 peptide sequences (1,259,215 with different modification types) were retained, which was 86.7% greater than what was initially reported by Kim *et al*. (293,700)^22^.

**Fig. 5.**
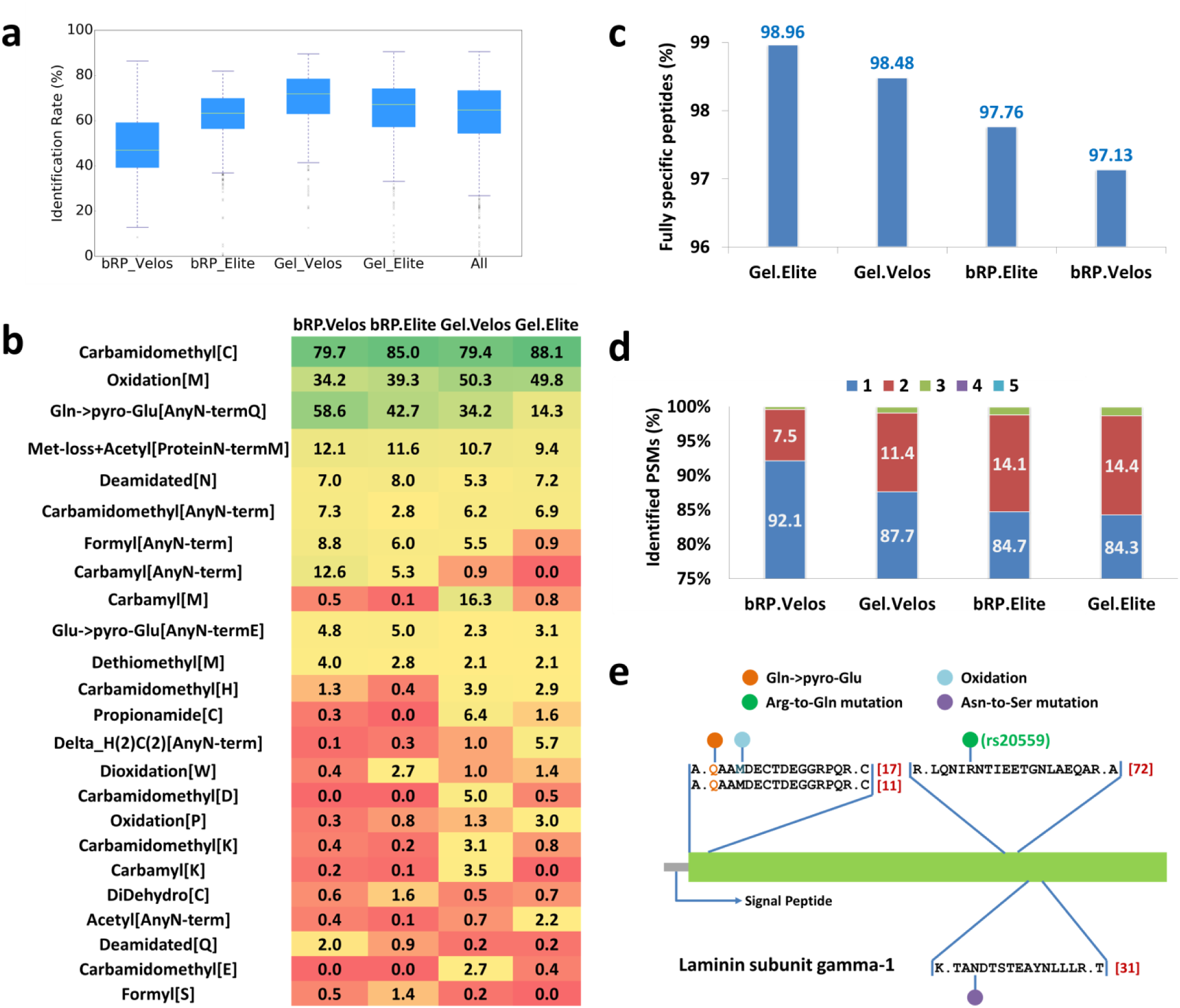
Overall analysis of the Kim data using Open-pFind. **a)** Distribution of the identification rate for each raw file. **b)** The distribution of highly abundant modifications discovered in the Kim data. Each number in one cell denotes the percentage of modified amino acids among all amino acids that appeared among the identified peptides. For example, 79.7% of cysteines were modified by carbamidomethylation in the identified peptides from an LTQ Orbitrap Velos MS fractionized by bRPLC. **c)** The proportions of fully specific peptides. **d)** Distribution of peptide numbers identified from one spectrum. For example, 7.5% of the identified spectra from an LTQ Orbitrap Velos MS fractionized by bRPLC contribute two peptides. **e)** The identified peptides in Laminin subunit gamma-1. Red numbers in the brackets denote how many PSMs correspond to each peptide.

The results obtained with Open-pFind demonstrated that the characteristics of MS/MS data vary according to different methods for sample preparation and LC-MS/MS. In terms of modifications, although several common modifications, *e.g.*, carbamidomethylation, oxidation and Gln→pyro-Glu, were always abundant in all datasets, many unexpected modifications still appeared in only one or two types of datasets (Fig. 5b). For example, propionamides of cysteines were hardly detected in the bRPLC fractionation samples but appeared in 1.6‒6.4% of all peptides from in-gel digested samples, which was consistent with a previous study by Sechi *et al*.^45^. On the other hand, the percentages of fully tryptic peptides were stable among the four types of datasets with different experimental conditions (97‒99% in Fig. 5c). In terms of co-eluting peptide identification, LTQ Orbitrap Elite tended to produce more mixed spectra than LTQ Orbitrap Velos, likely due to its higher sensitivity, allowing less-abundant peptides to be detected and identified via Open-pFind (Fig. 5d). The different characteristics of these datasets again proved that specifying an exact search space for each individual dataset based on expert experience is always difficult, and uniformly considering a complete search space for different experimental conditions is essential for today’s search engines.

Identification results from the extended search space were also valuable for biological discoveries. For example, a total of 9,559 semi-tryptic peptides were identified as being located in the N-terminal regions of proteins (the N-terminal amino acid of each peptide located between the 1^st^ and the 60^th^ amino acid of the corresponding protein), of which 34.1% had complete ion series (at least one *b* or *y* ion was detected at each peptide linkage), and 66.4% had at most two peptide linkages in which both the *b* and *y* ions were missing. These semi-tryptic peptides provide valuable clues for identifying signal peptides, and 694 of them were already verified in UniProt (Supplementary Table 10). The score distributions of these 9,559 peptides and the total 548,371 peptides (Supplementary Fig. 9) indicated that although these semi-tryptic peptides were from a much larger search space (Fig. 4b), their confidence was still comparable to that of the total results. On the other hand, biological modifications and mutations were effectively discovered by Open-pFind. For example, Laminin subunit gamma-1 was identified by different types of peptides, all of which were supported by over ten PSMs (Fig. 5e). The N-terminal cleavage site of QAAMDECTDEGGRPQR was confirmed by the signal peptide recorded in UniProt. In addition, two amino acid mutations were discovered by Open-pFind, and one of them, the R1121Q, was verified previously (rs20559 in dbSNP^46^).

Proteins were directly inferred by unique peptides that were not shared with any other proteins. The number of proteins supported by at least two unique peptides was 14,064, and the estimated FDR was 1% at the protein level, which corresponded to 12,723 genes (Fig. 6a). The average protein coverage was ∼41.5%, and for 11,500 proteins (81.8% of 14,064) the coverage was greater than 10% (Supplementary Table 11). On average, 38 peptides and 26 peptide sequences (regardless of modifications) were identified per protein. The number of proteins was also consistent with the statement by Ezkurdia *et al*. that the accurate gene number was approximately 12,000^23^. In addition, these authors performed a test that counted olfactory receptors, which should not be detected in standard proteomics experiments, and 108 olfactory receptors were found in the previously reported results. In contrast, if the two-peptide rule was used, the number of olfactory receptors reported by Open-pFind was two, which was only 0.016% of the total 14,064 proteins. A manual check further revealed that each of the two olfactory receptors was randomly matched with two peptides, each of which was supported by only one PSM. No olfactory receptors were found among the 12,239 proteins (11,536 genes) that were identified by three or more peptides. In summary, we believe that peptides and proteins are comprehensively and precisely identified via Open-pFind, even when analyzing an ultra-large proteome dataset.

**Fig. 6.**
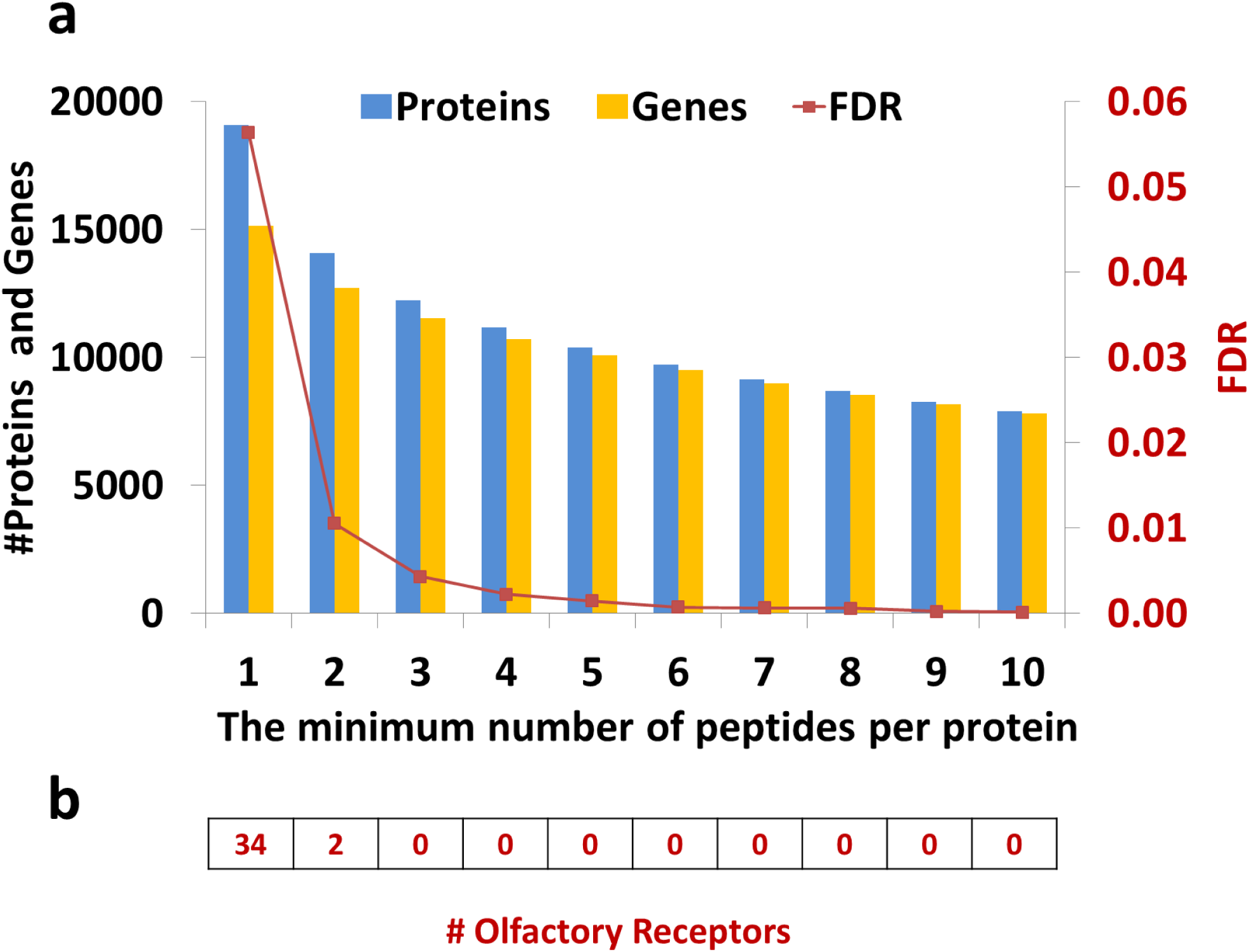
Protein and gene identification from the Kim data. **a)** Protein and gene numbers with different numbers of supporting peptides. **b)** The number of olfactory receptors with different numbers of supporting peptides.

## DISCUSSION

Thousands of chemical and biological modifications, as well as unexpected digestion types and mixed spectra, have led to low identification rates of MS/MS data. Therefore, an open search strategy, in which a more complete search space is considered, has become more important. Open-pFind has been proposed as an effective, accurate and fast open search algorithm, with the potential to be the most commonly used search engine for routine shotgun proteomics. In addition, the tag-based index approach and the two-step workflow are extensively applicable for many other types of search engines, *e.g.*, the identification of cross-linked peptides or glycopeptides. The open search strategy is also a quality control method to identify missing proteins and verify novel coding elements by easily providing competitive peptide identification based on the large search space^47^.

Protein inference, a separate problem downstream of peptide identification, was not extensively discussed in this study. Indeed, the protein inference strategies utilized by search engines are quite diverse and are difficult to evaluate comprehensively; thus, in this study, we used the simple but efficient two-peptide rule^25^. We also compared this rule with the ‘picked’ protein FDR approach^48^, designed to infer proteins in large-scale proteomic datasets, using the Kim data, and the performance of the two strategies was similar. The number of proteins reported by the picked strategy was 16,133, which was slightly larger than the number that was reported using the two-peptide rule (14,064), but seven additional olfactory receptors were detected with the picked strategy.

The metabolically labeled datasets proved to be effective when evaluating the accuracy of the identification results in this study. Additionally, these datasets may be further used to estimate the accuracy of the results, *i.e.*, the percentage of correct PSMs or peptides, based on two simple assumptions (Supplementary Note 4). In the Dong-Ecoli-QE dataset, the accuracy of the identified PSMs varied from 95.7‒99.2% for different engines when considering only the peptides in the restricted search space (Supplementary Fig. 10). For the separately identified results, the estimated accuracy of Open-pFind remained close to 99%, which was significantly higher in comparison with the other engines. Generally, if considering only peptides without any modifications or with only common modifications, all open search engines reported more precise results than those obtained with the restricted engines because the peptides from the restricted search space survived in a significantly larger space containing a huge number of competing peptide candidates. However, if all identified peptides were considered, the accuracy of the open search engines decreased to varying degrees. Open-pFind remained at a high global accuracy of 98.9%, while the accuracies of the other three open search engines dropped to 93.5% for the best, or to 86.6% for the worst. The potential of the metabolic labeling approach must be further explored. We suggest that quantitation information should be considered in designing the scoring model to improve search engine precision, especially for open search strategies, which will be increasingly popular for routine MS/MS data analyses in the future.

## ACKNOWLEDGMENTS

This work was supported by the National Key Research and Development Program of China (No. 2016YFA0501300, 2017YFA0505100, 2017YFC0906600), the CAS Interdisciplinary Innovation Team (Y604061000), the National Natural Science Foundation of China (31670834), the National Key Basic Research Program of China (No. 2012CB316502), the International Collaboration Program (2014DFB30020), Beijing Training Project for The Leading Talents in S&T (Z161100004916024), and the Youth Innovation Promotion Association CAS (No. 2014091).

## ONLINE METHODS

### Constructing the tag-index

For each protein in the specified database, all sequence tags with a specified length *k* are generated. For instance, given a protein sequence MAHVAEADK whose length is 9, the number of all its 3-mer tags is 7 (MAH, AHV, HVA, VAE, AEA, EAD and ADK). The tags extracted from all proteins are sorted according to the lexicographical order and then stored in a datasheet. For each tag, its protein ID (from 1 to *M* where *M* denotes the number of proteins in the database) and start position in the protein (from 1 to *L* where *L* denotes the length of the protein) are actually recorded rather than the real sequence, which can be compressed into only one 32-bit integer. Obviously, all tags are recorded in the datasheet with equal lengths, which is convenient for randomly retrieving any one of them. Finally, an index table is constructed, so that all occurrences of any one tag can be efficiently retrieved in protein databases. For each *k*-mer tag *T* with *k* amino acids *a*_1_ *a*_2…_*a_k_*, the key in the hash table is calculated by Formula 1, in which the function ascii is used to get the ASCII code of a given character. The time complexity of finding the first valid protein matched with the given tag in the datasheet is *O*(1).

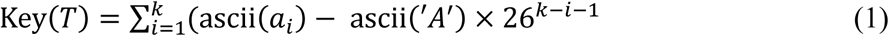

### Tag-index based open search

The workflow of the open search module, shown in Fig. 1b, adopted a tag-index to accelerate the retrieval of associated peptides. For each spectrum, a number of *k*-mer tags are extracted and then searched against the tag-indexed protein database. Generally, the parameter *k* is chosen as five to balance the search time (mainly correlated to the tag frequency, *i.e.*, the number of hits in a given database for each tag) and the storage of the index structure (Supplementary Table 12).

After finding the matched positions in the database, peptide candidates are generated by extending each of the matched tags to a full-length peptide sequence. Three types of extension procedures are supported in Open-pFind, resulting in three different open search strategies.

1. A maximum of one unexpected modification in the given modification list (*e.g.*, all modifications in Unimod^27^) is allowed for each peptide, which is the default search mode shown in this study. Peptides that fit at least one flanking mass of the tag are considered, and the mass shift on the other side is considered a potential modification if the mass appears in the given modification list. All modified peptides with different modification site localizations are generated and then scored for a given spectrum. For example, given the peptide AEHVASATK and phosphorylation as a potential modification, two modified peptides, AEHVA**pS**ATK and AEHVASA**pT**K, are generated, in which pS and pT denote the phosphorylation sites. Notably, there may be no valid modified forms generated for a given peptide due to the absence of any proper modification sites.
2. A maximum of two modifications in the given modification list are allowed for each peptide. First, the combinations of any two modifications are enumerated and stored in the memory. Each single modification is considered a special combination and stored together with the two-modification combinations. Second, all combinations are sorted in a list *C* according to their masses (the mass of one combination is computed by summing the masses of the modifications in it). Peptides that fit at least one flanking mass of the tag are considered, and the mass shift on the other side is considered to be contributed by a combination in *C*. Then, the combination is decoded, and all valid modified peptides are enumerated, which is similar though far more complex than the first strategy that allowed only one unexpected modification in each peptide.
3. Any masses within the given range are considered, *i.e.*, a blind search mode like that of MODa and MSFragger, in which no given modification lists are relied upon. Peptides that fit at least one flanking mass of the tag are considered, and the mass shift on the other side is considered a modification. Then, the modification is tested on all sites in the peptide (except those sites within the region matched with the tag) to generate different modified peptides in turn.

Finally, the peptide with the best score is chosen as the final result of the open search for each spectrum. The score function is the same as the previous versions of pFind. Generally, a few top peptide candidates (ten by default) are stored for each spectrum according to the settings determined by the users.

#### Reranking of PSMs

A semi-supervised learning algorithm is used to iteratively separate target PSMs from random PSMs based on the linear classification software package LIBLINEAR^49^. Six features are extracted from each PSM to train the scoring model, including 1) the original score, 2) the peptide length, 3) the ratio of the number of matched ions to that of all theoretical fragment ions, 4) the maximum tag length in the peptide, 5) the frequency of the specified modifications and 6) the frequency of the digestion type (fully, semi- or non-specific). The procedure to compute the last two features is the same as that previously reported by Chi *et al*^14^.

For each iteration, positive samples are formed by all target PSMs within the threshold of 1% FDR at the peptide level, and the negative samples are formed by all decoy PSMs out of the FDR threshold. Then, the model is trained and used to re-compute the score between one spectrum and each of its top peptide candidates. The candidates of each spectrum are sorted according to the new scores such that the top-ranked peptide may be changed in this step. Finally, new positive and negative samples are generated according to the FDR estimation for the new PSMs, and the next iteration is started. Generally, a maximum of ten iterations are needed to train a stable model.

The reranking module is used twice in the entire workflow of Open-pFind: after the open search and after the restricted search. For the second use, the results of each spectrum from both open and restricted searching are merged, sorted by their original scores and then reranked uniformly.

#### Refined search

Refined search starts after the reranking of the open search results. Generally, the module is designed the same as the restricted search engines, *e.g.*, pFind, MaxQuant and SEQUEST. However, there are two key differences between the restricted search module of Open-pFind and that of other traditional engines. First, after reranking, the protein database is automatically learned that only proteins supported by peptides within the specified FDR threshold (5% by default) are retained. The newly generated database is usually smaller than the original one, especially for those containing many irrelevant proteins, which will speed up the subsequent restricted search. In addition, given the sensitivity of the open search, nearly all target proteins in the experiment are considered in the restricted search. Second, variable modifications are determined according to the previous reranking step in which the frequency of each modification is computed. Generally, the top-*k* highly abundant modifications are selected (*k* is chosen as five by default). Relevant modifications are automatically and adaptively set using this strategy for different MS/MS datasets, unlike specifying modifications according to expert experience.

#### Mixed spectra analysis

A simple strategy is used to identify mixed spectra. First, *m* precursor ions are extracted using pParse (or other tools) for each spectrum. Secondly, *m* tandem mass spectra are generated by copying the MS/MS information of the original spectra and assigning *m* different precursor ions (monoisotopic masses and charge states) to *m* tandem mass spectra. Then, these newly generated spectra are searched in the traditional way by search engines.

However, this method may cause a serious problem. Given two spectra *A* and *B*, generated from the same original spectrum *S*, two peptides may be identified with an identical sequence but different modifications to fit the precursor ions of *A* and *B*. The two peptides may both match the spectrum well because many fragment ions are the same for the two peptides, and then match to the same peaks in *S* (although they may appear to have matched different peaks in *A* and *B*). However, if one peptide is considered the correct one, the other will be less credible due to the lack of independent evidence. The problem may be more serious for an open search procedure because more modifications are considered, resulting in a greater possibility that the mass gap between a precursor ion and the corresponding peptide will be randomly filled. In other words, given a mixed spectrum and several peptide candidates, the shared peaks matched with these peptides should be strictly limited to avoid reporting many incorrect PSMs with high scores.

Open-pFind improves the simple strategy described above at the beginning of this section by slightly modifying the reranking step. As shown in the reranking description, all spectra are sorted according to their scores matched with the top-ranked peptide before each iteration. For each PSM *p*, the improved strategy checks each of its siblings that were extracted from the original spectrum with the same scan number. If the score of a sibling PSM *q* is better than *p* and the number of shared peaks between these two PSMs *p* and *q* is greater than *k* (three by default), then *p* should be removed from the current PSM list (but it will still be considered in the next iteration because the scores of *p* and *q* may change). Generally, only the backbone ions are considered in this step, *e.g.*, *b* and *y* ions for HCD and *c* and *z* ions for ETD. In summary, the improved strategy guarantees that no two sibling PSMs from the same MS2 spectrum sharing more than *k* peaks are reported by Open-pFind; thus, most fragment ions in each retained PSM are uniquely matched in each mixed spectrum.

#### MS/MS datasets

First, two datasets were used to measure the sensitivity and accuracy of different engines. Dong-Ecoli-QE is derived from a sample of ^14^N-(*i.e.*, unlabeled), ^15^N- and ^13^C-labeled *E. coli* cultures at a ratio of 1:1:1, and Xu-Yeast-QEHF is derived from a sample of unlabeled and ^15^N-labeled yeast cultures at a ratio of 1:1. Then, another four published datasets obtained with different types of mass spectrometers were used to evaluate the performance of the search engines, *e.g.*, the MS/MS identification rate, the number of identified PSMs, peptides and proteins, and the consistency of the results between Open-pFind and other search engines. The details of these six datasets are shown in Supplementary Table 1.

#### Sample preparation for Dong-Ecoli-QE

The ^15^N-labeled *E. coli* cells were prepared as described^50^ using a M9 medium made of ^14^NH_4_Cl. The ^13^C-labeled *E. coli* cells were prepared in a similar way, except that the M9 medium was made of ^14^NH_4_Cl fully labeled ^13^C glucose. The bacterial cultures were grown for at least 24 h (eight generations) to complete ^15^N and ^13^C labeling. At OD_600_ around 0.8, the cells were harvested by centrifugation at 1000 × *g* and washed twice with a 10 mM Tris/HCl buffer (pH = 7). The bacterial cells were re-suspended in lysis buffer (4% SDS, 0.1 M Tris/HCl, pH 8.0), adjusted to OD_600_ of 7.5 per 100 μL, and disrupted by sonication for 10 min on ice. Unbroken cells were removed by centrifugation at 16,000 × *g* for 15 min. The protein concentration of the supernatant was determined using the bicinchoninic acid (BCA) method (Pierce), and the supernatant was stored at −80 °C.

#### Sample preparation for Xu-Yeast-QEHF

*Saccharomyces cerevisiae* SUB 592 was used for all experiments in this work. ^14^N and ^15^N labeling media were prepared by adding 0.1% (^15^NH_4_)_2_SO_4_ (99.14% atom percent excess, SRICI, Shanghai, China) or 0.1% (NH_4_)_2_SO_4_ to Synthetic Dextrose (SD) Medium (0.7% Difco yeast nitrogen base, 2% dextrose, supplemented with adenine and uracil) as described previously^51^. Seed cultures of SUB 592 were grown at 30°C with shaking (200 rpm) in ^14^N and ^15^N SD media (5 mL). The same cells (OD_600_, 0.05) were transferred in a 50-mL flask that included 10 mL of liquid ^14^N and ^15^N labeling media when minimal growth cultures were grown to mid-log phase. The ^14^N and ^15^N labeling cells were mixed 1:1 based on OD_600_ measurements when growing cells reached mid-log phase. The mixed labeling cells (8OD) were lysed in buffer (8 M urea, 5 mM IAA, 50 mM NH_4_HCO_3_, 1× protease cocktail) by the vortex mixer method (vortexed vigorously for 1 min, iced for 1 min, 10 cycles). The unbroken debris was eliminated by centrifugation (13,300×g) at 4°C for 10 min. The supernatant was collected and resolved by short SDS-PAGE (10%, 0.7 cm), followed by staining with Coomassie Brilliant Blue. The gel lanes were excised and digested with trypsin at 37°C for 14 h.

#### Database generation for Xu-Yeast-QEHF

The target protein database used for analyzing the Xu-Yeast-QEHF dataset contains three parts: 1) a six-frame translation database of the genome, 2) an N-terminal peptides database, and 3) a junction peptides database. We downloaded the whole genome sequence of yeast from SGD and then used a stop-codon-to-stop-codon six-frame translation strategy to the nuclear and mitochondria DNA. A standard codon table was used to translate nuclear DNA while a mitochondria codon table obtained from http://www.ncbi.nlm.nih.gov/Taxonomy/Utils/wprintgc.cgi was used to translate mitochondria DNA. ORFs containing less than six amino acids were removed from the database. We also list all the fully specific digestion peptides starting with methionine to build the N-terminal peptides database to ensure retrieving N-terminal peptides in a fully specific digestion search mode. In SGD, 284 coding genes had splice junctions and 47 coding genes had translational frame shifts. Junction peptides database included peptides spanning splice sites or translational frame shift sites meeting enzyme digestion rules. These peptides could guarantee the same phase position with the original annotated genes. In the end, we combined these three databases and common contaminants database and then reversed all the sequences to generate a decoy database to use the target-decoy identification strategy.

#### Database Search

Open-pFind and seven other search engines, specifically MSFragger, MODa, PEAKS-PTM (referred to as PEAKS), Comet, MS-GF+, Byonic and pFind, were investigated in this study. The open search mode was adopted in MSFragger, MODa and PEAKS, while the restricted search mode was used for the other four engines (Supplementary Table 2). To be more precise, PEAKS considered hundreds of modifications in its built-in modification list, while MODa and MSFragger employed a blind search mode that considered any mass shifts within a tolerance rather than the modifications pre-stored in a list such as Unimod. Non-tryptic peptides were considered in the search space for all open search engines with the exception of MSFragger because it always crashed when creating the ion index, even when using a server with 128 GB RAM. Therefore, only semi-tryptic peptides were searched by MSFragger across the six datasets for the running time comparison. In addition, the sensitivity of MSFragger was the highest when only tryptic peptides were considered (Supplementary Table 8); hence the results of MSFragger from the database search against fully tryptic peptides were used for the performance evaluation across the six datasets in the Results section. The FDR was controlled, when possible, to be 1% at the peptide level for the engines based on the target-decoy strategy with their primary scores (Open-pFind, MSFragger, Comet and pFind) or based on the built-in methods (MS-GF+, MODa, PEAKS and Byonic). For example, Byonic first controlled proteins at 1% FDR or at a maximum of 20 decoy hits and then estimated FDR at the spectrum level (generally 0-5%). All MS/MS data were analyzed using a standard desktop computer (8-core CPU @ 2.90 GHz and 32 GB RAM), in which six threads were specified for Open-pFind, MSFragger, pFind, Comet, MS-GF+ and Byonic (Multicore: Normal). MODa performed single-threaded searches because multiple threading was not supported in this version, and Open-pFind was also tested additionally with a single thread for a fair comparison. PEAKS used its built-in strategy (about 6–8 threads by observation from the task manager of the operating system).

#### Data availability

All raw files are described in Supplementary Table 1. The datasets of Dong-Ecoli-QE and Xu-Yeast-QEHF have been deposited into ProteomeXchange with the accession numbers of PXD008782 and PXD008783, respectively. The other four datasets were from the corresponding research articles published previously. Processed data files that support the findings of this study are available from the corresponding author upon request.

