## Supplementary Materials for "Open-pFind enables precise, comprehensive and rapid peptide identification in shotgun proteomics"

### Supplementary Tables 1-12

**Supplementary Table 1**. Detailed information on the six datasets used in this study

| Dataset | Mass spectrometer | # Raw files | # MS2 scans | Reference |
|---|---|---|---|---|
| **Dong-Ecoli-QE** | Q Exactive | 5 | 202,452 | / |
| **Xu-Yeast-QEHF** | Q Exactive HF | 22 | 526,301 | / |
| **Mann-Human-Velos** | LTQ Orbitrap Velos | 3 | 64,112 | [1] |
| **Gygi-Human-QE** | Q Exactive | 24 | 1,121,149 | [2] |
| **Mann-Mouse-QEHF** | Q Exactive HF | 4 | 746,116 | [3] |
| **Pandey-Human-Elite** [a] | LTQ Orbitrap Elite | 24 | 406,913 | [4] |

[a] Only the 24 raw files whose names begin with *Adult_CD8Tcells_Gel_Elite* were chosen.

**Supplementary Table 2.** The eight search engines used in this study

| Search engine | Version info. | Open search |
|---|---|---|
| **Open-pFind** | 1.0 | ✓ |
| **PEAKS** | 7.5 | ✓ |
| **MODa** | 1.23 | ✓ |
| **MSFragger** | v20170103 | ✓ |
| **pFind** | 3.1 | × |
| **Comet** | 2016012 | × |
| **MS-GF+** | v10072 | × |
| **Byonic** | 2.10 | × |

**Supplementary Table 3.** Parameters for database searches

| Items | Settings |
|---|---|
| **Database** | Target + Decoy [a] |
| **Enzyme** | Trypsin |
| **Digestion** | Fully specific for restricted engines and MSFragger<br>Non-Specific for Open-pFind, MODa and PEAKS |
| **Max. missed cleavage sites** | 3 |
| **Mass tolerance of precursor ions** | ± 20 ppm |
| **Mass tolerance of fragment ions** | ± 20 ppm (± 0.02 Da if the ppm unit is not supported for the search engines, *e.g.*, PEAKS) |
| **Modifications** | Fixed: carbamidomethylation (C)<br>Variable: oxidation (M), Gln→pyro-Glu (N-termini of peptides) and acetyl (N-termini of proteins) |

[a] The human protein database was downloaded from UniProt (2016-4-20) for Mann-Human-Velos, Gygi-Human-QE and Pandey-Human-Elite. The mouse protein database was downloaded from UniProt (2016-11-29) for Mann-Mouse-QEHF. Both reviewed and unreviewed proteins were used in this study by default. The *E. coli* protein database for the K-12 substrain MG1655 downloaded from NCBI on 2015-10-14 was used for Dong-Ecoli-QE. The six-frame-translated database was used as the target database for Xu-Yeast-QEHF (Online Methods).

**Supplementary Table 4.** Consistency of the identification results

(separate Excel file)

**Supplementary Table 5.** The average number of peptides per protein in the consistently identified proteins in the Dong-Ecoli-QE dataset

| Search engine | # Peptides per protein |
|---|---|
| **Open-pFind** | 17.3 |
| **PEAKS** | 15.3 |
| **MSFragger** | 14.0 |
| **MODa** | 10.1 |
| **Byonic** | 9.6 |
| **pFind** | 9.2 |
| **Comet** | 9.0 |
| **MS-GF+** | 8.9 |

**Supplementary Table 6.** Classification of the revived spectra from different engines in the Dong-Ecoli-QE dataset

| Search engine | # Non-nested peptides | # Nested peptides | # Nested peptides / # total peptides (%) |
|---|---|---|---|
| **MSFragger** | 492 | 3,669 | 88.2 |
| **PEAKS** | 130 | 1,091 | 89.4 |
| **MODa** | 327 | 2,950 | 90.0 |
| **pFind** | 15 | 25 | 62.5 |
| **MS-GF+** | 41 | 770 | 94.9 |
| **Comet** | 49 | 807 | 94.3 |
| **Byonic** | 176 | 21 | 10.7 |

**Supplementary Table 7.** Real search times (in minutes) of the eight search engines for the six datasets

| | pFind | Byonic | MS-GF+ | Comet | MSFragger | PEAKS | MODa | Open-pFind [a] |
|---|---|---|---|---|---|---|---|---|
| **Xu-Yeast-QEHF** | 73 | 144 | 85 | 118 | 5071 | 1,269 | 4,896 | 41 (158) |
| **Dong-Ecoli-QE** | 4 | 36 | 41 | 22 | 130 | 448 | 91 | 8 (32) |
| **Mann-Human-Velos** | 9 | 20 | 19 | 26 | 747 | 167 | 1291 | 20 (78) |
| **Mann-Mouse-QEHF** | 101 | 274 | 605 | 623 | 6260 | 12,178 | 52,228 | 263 (1,210) |
| **Gygi-Human-QE** | 94 | 347 | 383 | 664 | 12137 | 17,013 | 27,469 | 210 (903) |
| **Pandey-Human-Elite** | 61 | 135 | 186 | 483 | 4368 | 5,880 | 12,440 | 92 (414) |

Note: All MS/MS data were analyzed using a standard desktop computer (8-core CPU @ 2.90 GHz and 32 GB RAM), in which six threads were specified for Open-pFind, MSFragger, pFind, Comet, MS-GF+ and Byonic (Multicore: Normal). MODa performed single-thread searches because multiple threading was not supported in this version. PEAKS used its built-in strategy (about 6–8 threads by observation from the task manager of the operating system).

[a] The single-threaded search time is shown in parentheses.

**Supplementary Table 8.** The analysis of a single LC-MS/MS run consisting of 41,820 MS/MS spectra in the Gygi-Human-QE dataset

| | Fully Specific | | Semi-Specific | | Non-Specific | |
|---|---|---|---|---|---|---|
| | Time | # PSM | Time | # PSM | Time | # PSM |
| **MODa** | 136 | 14,370 | 179 | 19,593 | 249 | 19,748 |
| **PEAKS** | 123 | 23,578 | 164 | 26,300 | 324 | 26,194 |
| **MSFragger** | 16 | 22,768 | 453 | 20,239 | 2,466 | 18,898 |
| **Open-pFind** | 8 | 36,369 | 9 | 37,895 | 9 | 37,854 |
| **Open-pFind (Unimod-2)** | 12 | 35,929 | 21 | 37,577 | 31 | 37,487 |
| **Open-pFind (Blind)** | 7 | 36,075 | 11 | 36,421 | 18 | 36,304 |

Note: The running time is measured in minutes.

**Supplementary Table 9.** The results of three open search engines with the *T. tengcongensis* dataset

| | Fully specific | | Non-specific | |
|---|---|---|---|---|
| | Time (min.) | # PSM | Time (min.) | # PSM |
| **PEAKS** | 205 | 40,850 | 268 | 69,521 |
| **MODa** | 33 | 26,084 | 60 | 35,941 |
| **MSFragger** | 8 | 38,004 | 291 | 44,794 |
| **Open-pFind** | 4 | 48,564 | 6 | 70,829 |

Note: The dataset contains 113,531 tandem mass spectra. The *T. tengcongensis* database was downloaded from UniProt (2017-05-04), containing both reviewed and unreviewed proteins. The other parameters were the same as those for the other analyses in this study.

**Supplementary Table 10.** Detailed information for 694 semi-tryptic peptides verified by UniProt (separate Excel file)

**Supplementary Table 11.** The number of identified proteins and genes on Kim data

| Min. Pep. | Proteins | FDR (%) | Genes | Olfactory receptor | Average coverage (%) | | Low coverage (< 10%) proteins | |
|---|---|---|---|---|---|---|---|---|
| | | | | | All pep. | Distinct pep. | All pep. | Distinct pep. |
| **1** | 19,067 | 5.63 | 15,153 | 34 | 32.0 | 27.9 | 7,282 | 8,762 |
| **2** | 14,064 | 1.05 | 12,723 | 2 | 41.5 | 36.5 | 2,564 | 3,608 |
| **3** | 12,239 | 0.43 | 11,536 | 0 | 46.4 | 41.2 | 1,231 | 1,948 |
| **4** | 11,168 | 0.22 | 10,708 | 0 | 49.5 | 44.3 | 707 | 1184 |
| **5** | 10,387 | 0.14 | 10,069 | 0 | 51.8 | 46.7 | 436 | 775 |
| **6** | 9,718 | 0.07 | 9,494 | 0 | 53.7 | 48.6 | 276 | 525 |
| **7** | 9,148 | 0.07 | 8,980 | 0 | 55.3 | 50.2 | 188 | 367 |
| **8** | 8,682 | 0.06 | 8,549 | 0 | 56.7 | 51.7 | 132 | 268 |
| **9** | 8,273 | 0.02 | 8,162 | 0 | 57.9 | 53.0 | 93 | 194 |
| **10** | 7,899 | 0.01 | 7,799 | 0 | 59.1 | 54.1 | 69 | 146 |

**Supplementary Table 12.** The relationship between the tag length and the performance of tag-index [a]

| Tag length | Average frequency | Storage space (MB) |
|---|---|---|
| **2** | 88621.2 | 3.1E-03 |
| **3** | 4954.3 | 6.1E-02 |
| **4** | 273.6 | 1.2 |
| **5** | 19.0 | 24.4 |
| **6** | 5.2 | 488.3 |

[a] Reviewed and unreviewed human proteins (152,493 in total) were downloaded from UniProt and used in this study.
