## Supplementary Materials for "Open-pFind enables precise, comprehensive and rapid peptide identification in shotgun proteomics"

### Supplementary Figures 1-10

**Supplementary Fig. 1.** Proportions of the NaN-ratio PSMs identified in the Dong-Ecoli-QE dataset. ... 1

**a** $^{15}N/^{14}N$: NaN Ratio of the consistently identified PSMs (%)

| Row. ∩ Col. | Open-pFind | MSFragger | PEAKS | MODa | pFind | MS-GF+ | Comet | Byonic |
|---|---|---|---|---|---|---|---|---|
| Open-pFind | 0.67 | | | | | | | |
| MSFragger | 0.21 | 1.32 | | | | | | |
| PEAKS | 0.63 | 0.24 | 1.17 | | | | | |
| MODa | 0.17 | 0.83 | 0.20 | 1.05 | | | | |
| pFind | 0.55 | 0.18 | 0.52 | 0.16 | 0.65 | | | |
| MS-GF+ | 0.52 | 0.67 | 0.61 | 0.65 | 0.52 | 2.91 | | |
| Comet | 0.25 | 0.77 | 0.35 | 0.73 | 0.24 | 2.10 | 2.76 | |
| Byonic | 0.45 | 0.20 | 0.50 | 0.18 | 0.46 | 0.48 | 0.22 | 1.77 |

**b** $^{15}N/^{14}N$: NaN Ratio of the separately identified PSMs (%)

| Row. – Col. | Open-pFind | MSFragger | PEAKS | MODa | pFind | MS-GF+ | Comet | Byonic |
|---|---|---|---|---|---|---|---|---|
| Open-pFind | | 1.05 | 0.72 | 1.02 | 0.81 | 0.83 | 1.13 | 0.87 |
| MSFragger | 9.61 | | 4.80 | 2.51 | 4.16 | 3.20 | 3.08 | 4.44 |
| PEAKS | 5.98 | 2.53 | | 2.55 | 3.74 | 2.93 | 4.09 | 3.09 |
| MODa | 12.03 | 2.11 | 6.65 | | 4.33 | 2.78 | 2.60 | 4.72 |
| pFind | 2.82 | 1.46 | 2.04 | 1.43 | | 1.34 | 3.40 | 1.43 |
| MS-GF+ | 23.72 | 7.51 | 20.57 | 6.94 | 17.50 | | 14.65 | 12.07 |
| Comet | 21.89 | 6.69 | 18.07 | 6.12 | 16.07 | 8.06 | | 11.45 |
| Byonic | 11.37 | 5.27 | 12.36 | 4.88 | 9.21 | 7.62 | 10.06 | |

**c** $^{13}C/^{12}C$: NaN Ratio of the consistently identified PSMs (%)

| Row. ∩ Col. | Open-pFind | MSFragger | PEAKS | MODa | pFind | MS-GF+ | Comet | Byonic |
|---|---|---|---|---|---|---|---|---|
| Open-pFind | 0.91 | | | | | | | |
| MSFragger | 0.31 | 1.41 | | | | | | |
| PEAKS | 0.84 | 0.33 | 1.62 | | | | | |
| MODa | 0.22 | 0.89 | 0.27 | 1.19 | | | | |
| pFind | 0.81 | 0.30 | 0.74 | 0.23 | 1.03 | | | |
| MS-GF+ | 0.75 | 0.78 | 0.83 | 0.72 | 0.76 | 3.28 | | |
| Comet | 0.44 | 0.89 | 0.51 | 0.82 | 0.44 | 2.22 | 3.11 | |
| Byonic | 0.65 | 0.30 | 0.71 | 0.25 | 0.70 | 0.69 | 0.43 | 3.03 |

**d** $^{13}C/^{12}C$: NaN Ratio of the separately identified PSMs (%)

| Row. – Col. | Open-pFind | MSFragger | PEAKS | MODa | pFind | MS-GF+ | Comet | Byonic |
|---|---|---|---|---|---|---|---|---|
| Open-pFind | | 1.42 | 1.00 | 1.40 | 1.04 | 1.08 | 1.44 | 1.15 |
| MSFragger | 9.72 | | 4.86 | 2.66 | 4.18 | 3.26 | 3.12 | 4.52 |
| PEAKS | 8.74 | 3.53 | | 3.58 | 5.16 | 4.18 | 5.66 | 4.26 |
| MODa | 13.28 | 2.58 | 7.22 | | 4.68 | 3.17 | 2.97 | 5.13 |
| pFind | 5.97 | 2.30 | 3.98 | 2.28 | | 2.48 | 4.93 | 2.40 |
| MS-GF+ | 25.39 | 8.46 | 22.16 | 7.87 | 18.77 | | 18.89 | 13.07 |
| Comet | 23.42 | 7.48 | 19.55 | 6.89 | 17.17 | 10.31 | | 12.27 |
| Byonic | 20.32 | 9.11 | 22.22 | 8.46 | 16.25 | 13.57 | 16.94 | |

**Supplementary Fig. 1.** Proportions of the NaN-ratio PSMs identified in the Dong-Ecoli-QE dataset. For the open search engines, only PSMs without any modifications or with modifications specified in the restricted search engines were considered in the statistical analysis.

**a)** Consistently identified PSMs from the comparison of every two search engines are considered.

**b)** Separately identified PSMs from the comparison of every two search engines are considered. The $^{15}N$- and unlabeled peptides are used to calculate the quantitative values in both a) and b).

**c)** Similar to a), but the $^{13}C$- and unlabeled peptides are used to calculate the quantitative values.

**d)** Similar to b), but the $^{13}C$- and unlabeled peptides are used to calculate the quantitative values.

**a**

$^{15}N$ / $^{14}N$: NaN Ratio of the consistently identified PSMs (%)

| Row. ∩ Col. | Open-pFind | MSFragger | PEAKS | MODa | pFind | MS-GF+ | Comet | Byonic |
|---|---|---|---|---|---|---|---|---|
| Open-pFind | 1.26 | | | | | | | |
| MSFragger | 0.42 | 13.31 | | | | | | |
| PEAKS | 0.90 | 1.17 | 4.33 | | | | | |
| MODa | 0.31 | 4.56 | 1.19 | 10.17 | | | | |
| pFind | 0.55 | 0.21 | 0.53 | 0.16 | 0.65 | | | |
| MS-GF+ | 0.61 | 0.75 | 0.68 | 0.65 | 0.52 | 2.91 | | |
| Comet | 0.32 | 0.93 | 0.40 | 0.82 | 0.24 | 2.10 | 2.76 | |
| Byonic | 0.45 | 0.20 | 0.50 | 0.18 | 0.46 | 0.48 | 0.22 | 1.77 |

**b**

$^{15}N$ / $^{14}N$: NaN Ratio of the separately identified PSMs (%)

| Row. – Col. | Open-pFind | MSFragger | PEAKS | MODa | pFind | MS-GF+ | Comet | Byonic |
|---|---|---|---|---|---|---|---|---|
| Open-pFind | | 2.11 | 1.74 | 2.02 | 1.87 | 1.77 | 2.06 | 1.80 |
| MSFragger | 48.02 | | 33.14 | 26.87 | 28.37 | 30.35 | 31.02 | 26.23 |
| PEAKS | 20.31 | 8.42 | | 8.31 | 11.66 | 11.08 | 11.98 | 10.12 |
| MODa | 54.18 | 26.91 | 39.07 | | 26.75 | 30.10 | 30.89 | 24.25 |
| pFind | 3.00 | 1.87 | 2.12 | 1.67 | | 1.34 | 3.40 | 1.43 |
| MS-GF+ | 11.26 | 7.08 | 10.77 | 5.25 | 17.50 | | 14.65 | 12.07 |
| Comet | 10.91 | 5.43 | 10.06 | 4.17 | 16.07 | 8.06 | | 11.45 |
| Byonic | 11.45 | 5.57 | 12.67 | 4.87 | 9.21 | 7.62 | 10.06 | |

**c**

$^{13}C$ / $^{12}C$: NaN Ratio of the consistently identified PSMs (%)

| Row. ∩ Col. | Open-pFind | MSFragger | PEAKS | MODa | pFind | MS-GF+ | Comet | Byonic |
|---|---|---|---|---|---|---|---|---|
| Open-pFind | 1.90 | | | | | | | |
| MSFragger | 0.78 | 15.54 | | | | | | |
| PEAKS | 1.25 | 1.45 | 5.00 | | | | | |
| MODa | 0.64 | 6.21 | 1.44 | 11.72 | | | | |
| pFind | 0.80 | 0.34 | 0.76 | 0.28 | 1.03 | | | |
| MS-GF+ | 0.85 | 0.85 | 0.91 | 0.73 | 0.76 | 3.28 | | |
| Comet | 0.51 | 1.02 | 0.56 | 0.90 | 0.44 | 2.22 | 3.11 | |
| Byonic | 0.65 | 0.31 | 0.72 | 0.25 | 0.70 | 0.69 | 0.43 | 3.03 |

**d**

$^{13}C$ / $^{12}C$: NaN Ratio of the separately identified PSMs (%)

| Row. – Col. | Open-pFind | MSFragger | PEAKS | MODa | pFind | MS-GF+ | Comet | Byonic |
|---|---|---|---|---|---|---|---|---|
| Open-pFind | | 2.94 | 2.57 | 2.84 | 2.83 | 2.52 | 2.86 | 2.72 |
| MSFragger | 53.04 | | 36.84 | 30.34 | 31.48 | 33.31 | 34.01 | 30.68 |
| PEAKS | 23.16 | 9.77 | | 9.66 | 13.09 | 12.26 | 13.37 | 11.54 |
| MODa | 56.95 | 28.25 | 41.66 | | 28.52 | 31.60 | 32.34 | 27.87 |
| pFind | 6.35 | 2.93 | 4.28 | 2.62 | | 2.48 | 4.93 | 2.40 |
| MS-GF+ | 15.47 | 8.71 | 14.37 | 6.60 | 18.77 | | 18.89 | 13.07 |
| Comet | 14.70 | 6.83 | 13.28 | 5.33 | 17.17 | 10.31 | | 12.27 |
| Byonic | 20.52 | 9.64 | 22.84 | 8.45 | 16.25 | 13.57 | 16.94 | |

**Supplementary Fig. 2.** Proportions of NaN-ratio PSMs identified in the Dong-Ecoli-QE dataset. All PSMs were considered in the statistical analysis.

**a)** Consistently identified PSMs from the comparison of every two search engines are considered.

**b)** Separately identified PSMs from the comparison of every two search engines are considered. The $^{15}N$- and unlabeled peptides are used to calculate the quantitative values in both a) and b).

**c)** Similar to a), but the $^{13}C$- and unlabeled peptides are used to calculate the quantitative values.

**d)** Similar to b), but the $^{13}C$- and unlabeled peptides are used to calculate the quantitative values.

**a** $^{15}N$ / $^{14}N$: NaN Ratio of the consistently identified PSMs (%)

| Row. ∩ Col. | Open-pFind | MSFragger | PEAKS | MODa | pFind | MS-GF+ | Comet | Byonic |
|---|---|---|---|---|---|---|---|---|
| Open-pFind | 0.45 | | | | | | | |
| MSFragger | 0.32 | 1.32 | | | | | | |
| PEAKS | 0.31 | 0.36 | 0.63 | | | | | |
| MODa | 0.07 | 0.60 | 0.10 | 0.89 | | | | |
| pFind | 0.32 | 0.33 | 0.31 | 0.08 | 0.37 | | | |
| MS-GF+ | 0.34 | 0.95 | 0.43 | 0.48 | 0.35 | 2.97 | | |
| Comet | 0.08 | 0.82 | 0.20 | 0.58 | 0.10 | 2.05 | 2.79 | |
| Byonic | 0.30 | 0.33 | 0.35 | 0.08 | 0.32 | 0.36 | 0.09 | 2.62 |

**b** $^{15}N$ / $^{14}N$: NaN Ratio of the separately identified PSMs (%)

| Row. – Col. | Open-pFind | MSFragger | PEAKS | MODa | pFind | MS-GF+ | Comet | Byonic |
|---|---|---|---|---|---|---|---|---|
| Open-pFind | | 0.53 | 0.73 | 0.71 | 0.67 | 0.58 | 0.81 | 0.64 |
| MSFragger | 10.56 | | 7.82 | 4.01 | 5.98 | 3.65 | 3.31 | 6.57 |
| PEAKS | 2.79 | 0.89 | | 1.22 | 1.88 | 1.13 | 1.39 | 1.48 |
| MODa | 10.66 | 1.91 | 10.13 | | 5.63 | 3.87 | 2.11 | 6.97 |
| pFind | 1.10 | 0.42 | 1.02 | 0.75 | | 0.43 | 1.02 | 0.55 |
| MS-GF+ | 21.08 | 5.91 | 21.55 | 6.92 | 17.29 | | 6.69 | 10.92 |
| Comet | 20.40 | 5.94 | 20.21 | 6.38 | 16.22 | 8.96 | | 10.36 |
| Byonic | 12.32 | 5.15 | 14.28 | 5.85 | 9.82 | 7.53 | 6.44 | |

**Supplementary Fig. 3.** Proportions of the NaN-ratio PSMs identified in the Xu-Yeast-QEHF dataset. For the open search engines, only PSMs without any modifications or with modifications specified in the restricted search engines were considered in the statistical analysis.

**a)** Consistently identified PSMs from the comparison of every two search engines are considered.

**b)** Separately identified PSMs from the comparison of every two search engines are considered.

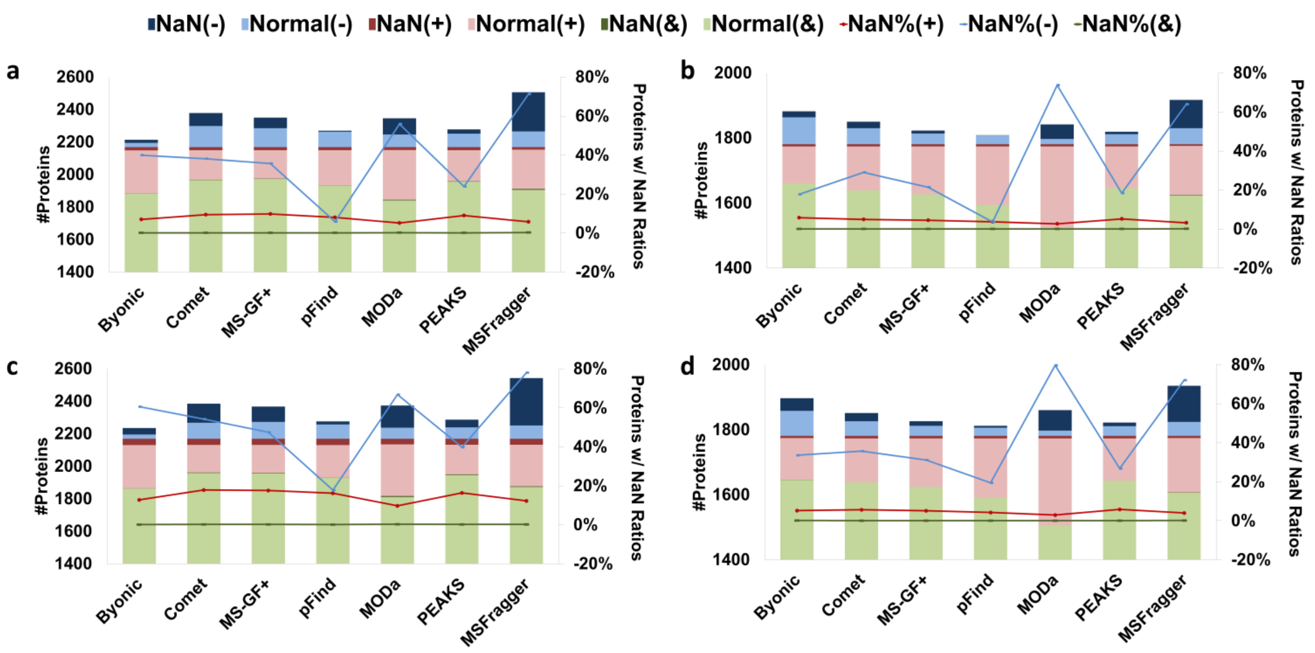

**Supplementary Fig. 4.** Analysis of NaN-ratio proteins in the Dong-Ecoli-QE dataset. Open-pFind is compared with each of the other seven engines.

**a)** The number of proteins that are consistently identified by two engines (&), separately identified by Open-pFind (+) or the other engine (–), and the proportions of NaN-ratio proteins in those three sets (&, + and –, respectively). The quantitative value of a protein is the NaN ratio if there are no supporting peptides with normal quantitative values.

**b)** Similar to a), but proteins supported with at least two peptides are considered. The quantitative value of a protein is the NaN ratio if the number of the supporting peptides with normal quantitative values is less than two.

$^{15}N$-labeled peptides were used to calculate the quantitative values in both a) and b).

**c)** Similar to a), but $^{13}C$-labeling peptides were used for calculating the quantitative values.

**d)** Similar to b), but $^{13}C$-labeling peptides were used for calculating the quantitative values.

**a**

| | Open-pFind | MSFragger | PEAKS | MODa | pFind | MS-GF+ | Comet | Byonic |
|---|---|---|---|---|---|---|---|---|
| Open-pFind | | 51.4 | 22.7 | 48.9 | 13.8 | 39.0 | 54.7 | 51.0 |
| MSFragger | 9.5 | | 22.8 | 4.5 | 19.4 | 11.5 | 8.8 | 38.4 |
| PEAKS | 1.5 | 15.5 | | 20.0 | 1.9 | 9.1 | 14.6 | 21.0 |
| MODa | 9.8 | 8.3 | 25.0 | | 19.2 | 7.4 | 8.9 | 44.0 |
| pFind | 1.4 | 25.4 | 7.7 | 31.1 | | 11.1 | 24.2 | 17.3 |
| MS-GF+ | 4.8 | 7.7 | 6.1 | 8.8 | 3.0 | | 0.8 | 16.8 |
| Comet | 8.9 | 9.9 | 14.0 | 7.7 | 9.9 | 1.4 | | 34.7 |
| Byonic | 3.8 | 20.8 | 6.8 | 28.2 | 3.6 | 22.3 | 36.8 | |

**b**

| | Open-pFind | MSFragger | PEAKS | MODa | pFind | MS-GF+ | Comet | Byonic |
|---|---|---|---|---|---|---|---|---|
| Open-pFind | | 93.8 | 71.8 | 95.5 | 15.9 | 77.8 | 87.1 | 55.3 |
| MSFragger | 20.2 | | 45.8 | 43.2 | 22.2 | 60.8 | 54.9 | 54.0 |
| PEAKS | 8.2 | 74.4 | | 76.7 | 5.2 | 72.0 | 80.8 | 27.5 |
| MODa | 23.2 | 34.3 | 42.4 | | 24.3 | 67.7 | 63.1 | 54.3 |
| pFind | 4.5 | 92.2 | 51.0 | 95.0 | | 11.1 | 24.2 | 17.3 |
| MS-GF+ | 9.8 | 72.9 | 47.2 | 79.9 | 3.0 | | 0.8 | 16.8 |
| Comet | 22.6 | 71.8 | 66.0 | 77.4 | 9.9 | 1.4 | | 34.7 |
| Byonic | 10.2 | 87.2 | 48.6 | 91.9 | 3.6 | 22.3 | 36.8 | |

**Supplementary Fig. 5.** The proportions of NaN-ratio PSMs obtained from one engine but revived by others.

**a)** Comparison between every two search engines. Each decimal denotes the percentage of PSMs revived by the search engine in the row (leftmost) for the total NaN-ratio PSMs from the search engine in the column (topmost). Only peptides with common modifications are considered.

**b)** Similar to a), but all PSMs including all types of modifications are considered. The $^{15}N$-labeled peptides and the unlabeled (common) peptides are used to calculate the quantitative values.

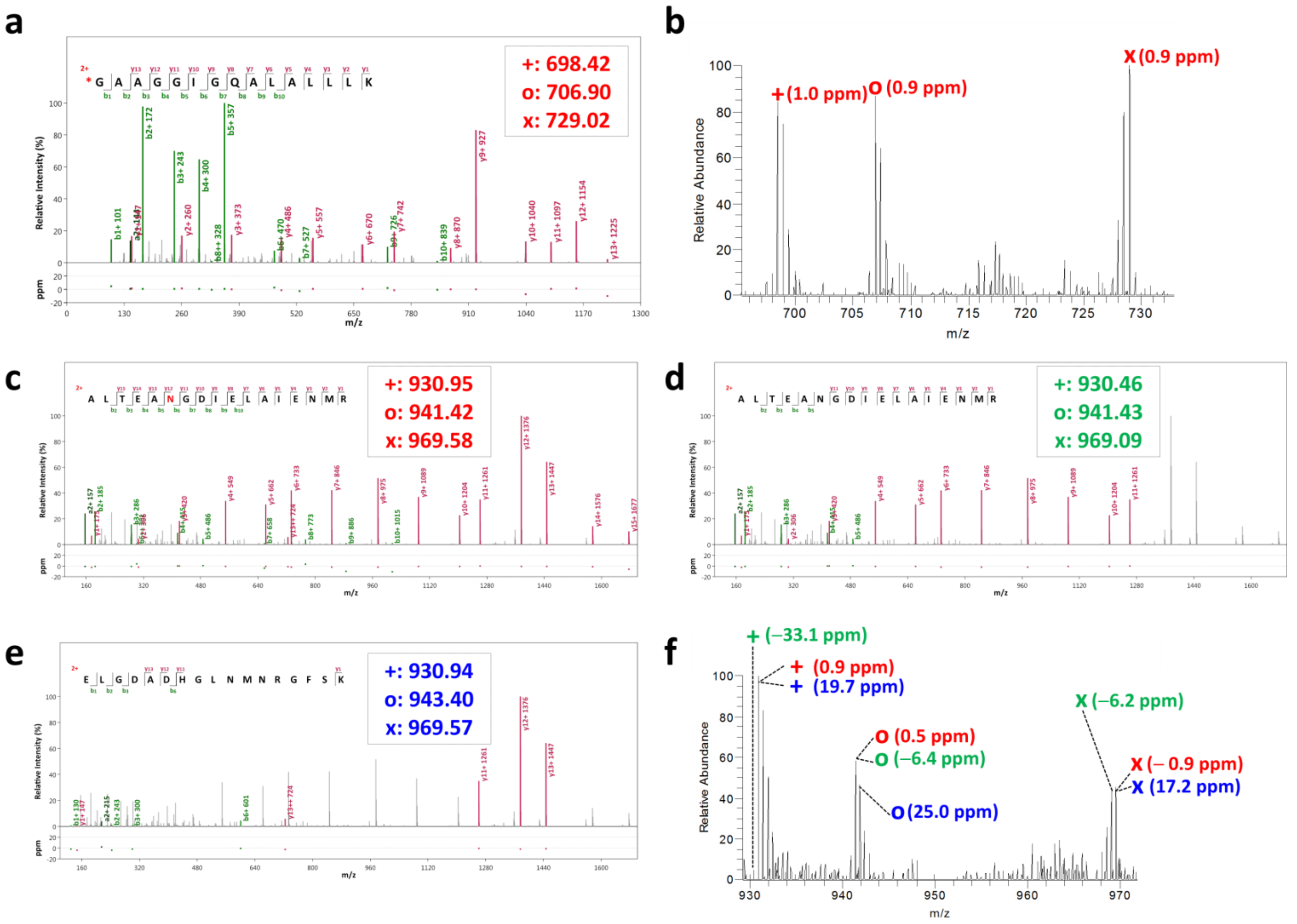

**Supplementary Fig. 6.** Two example spectra showing the effects of the metabolic labeling technique to distinguish the correct PSMs. +, o and x denote the monoisotopic masses of the unlabeled, 15N- and 13C-labeled precursor ions, respectively.

- **a)** Ecoli-1to1to1-un-C13-N15-60mM-20150823.42526.42526.2.dta, which is identified by Open-pFind as a semi-tryptic peptide, GAAGGIGQALALLLK, with a carbamylation at the N-terminus (m/z = 698.4203). MSFragger reported another peptide, ***VAVL***GAAGGLGQALALLLK (m/z = 699.3906, Hyperscore= 13.5427). If the precursor ion m/z was changed to 698.4203 for MSFragger (the same to that used in Open-pFind) and semi-tryptic peptides were allowed to search against, a new peptide GAAGGLGQALALLLK was reported with a mass shift of 43.0074 Da (The monoisotopic mass of carbamylation), whose Hyperscore was 35.7024.
- **b)** The MS1 information corresponding to the PSM shown in a).
- **c)** Ecoli-1to1to1-un-C13-N15-30mM-20150823.35791.35791.2.dta, which is identified by Open-pFind as a peptide, ALTEANGDIELAIENMR, with a deamidation of N at the 6th position.
- **d)** The same spectrum as c), which is identified by Comet and MS-GF+ as a peptide, ALTEANGDIELAIENMR, without any modifications.
- **e)** The same spectrum as c), which is identified by Byonic as a peptide, ELGDADHGLNMNRGFSK, without any modifications.
- **f)** The MS1 information corresponding to the PSMs shown in c)-e).

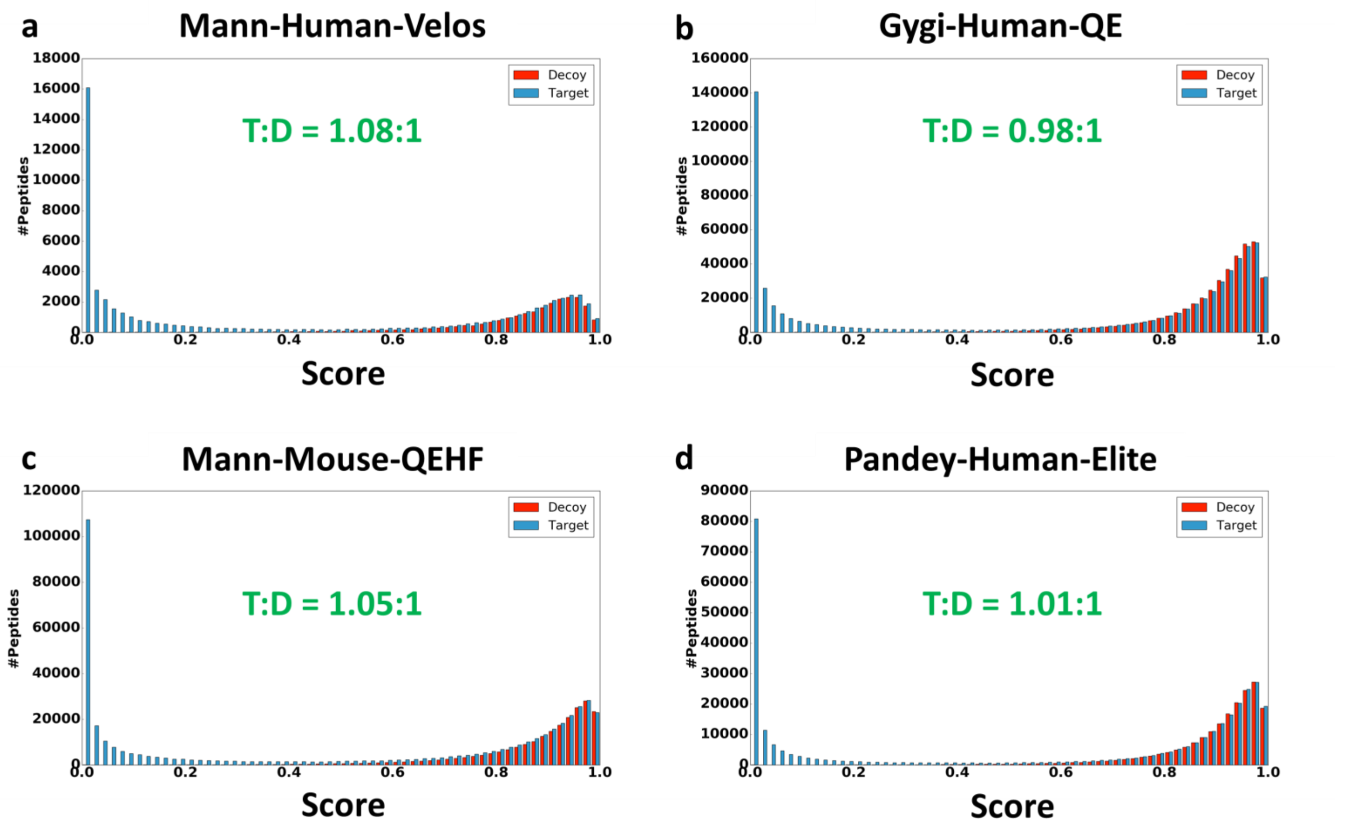

**Supplementary Fig. 7.** The score distribution of the target (blue) and decoy (red) peptides identified by Open-pFind from four published datasets: **a)** Mann-Human-Velos, **b)** Gygi-Human-QE, **c)** Mann-Mouse-QEHF and **d)** Pandey-Human-Elite. The range of the scores is 0–1, and a larger score value indicates the better quality of one PSM. All results, without any FDR control, are shown. The ratio shown in each figure was computed using the peptides with scores higher than 0.8.

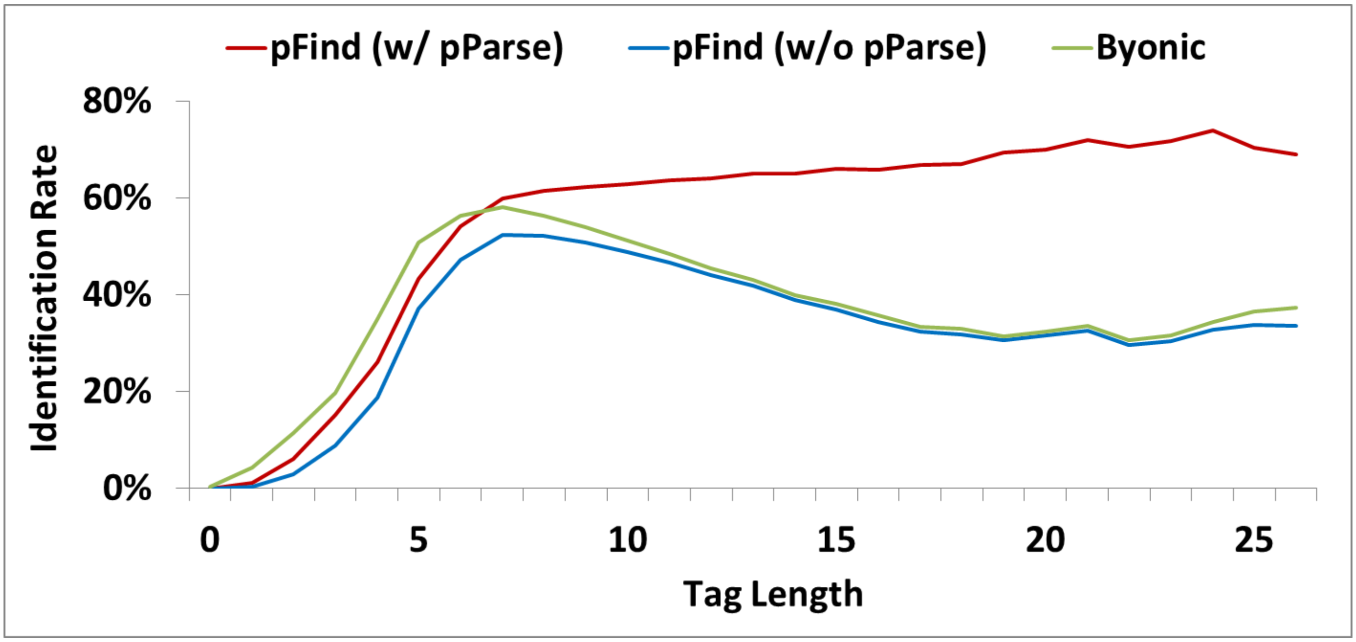

**Supplementary Fig. 8.** The distribution of the identification rates of Byonic and pFind at different maximum tag lengths extracted from the spectra. Two modes are adopted for pFind, and the only difference is whether pParse is used to calibrate the precursor ions.

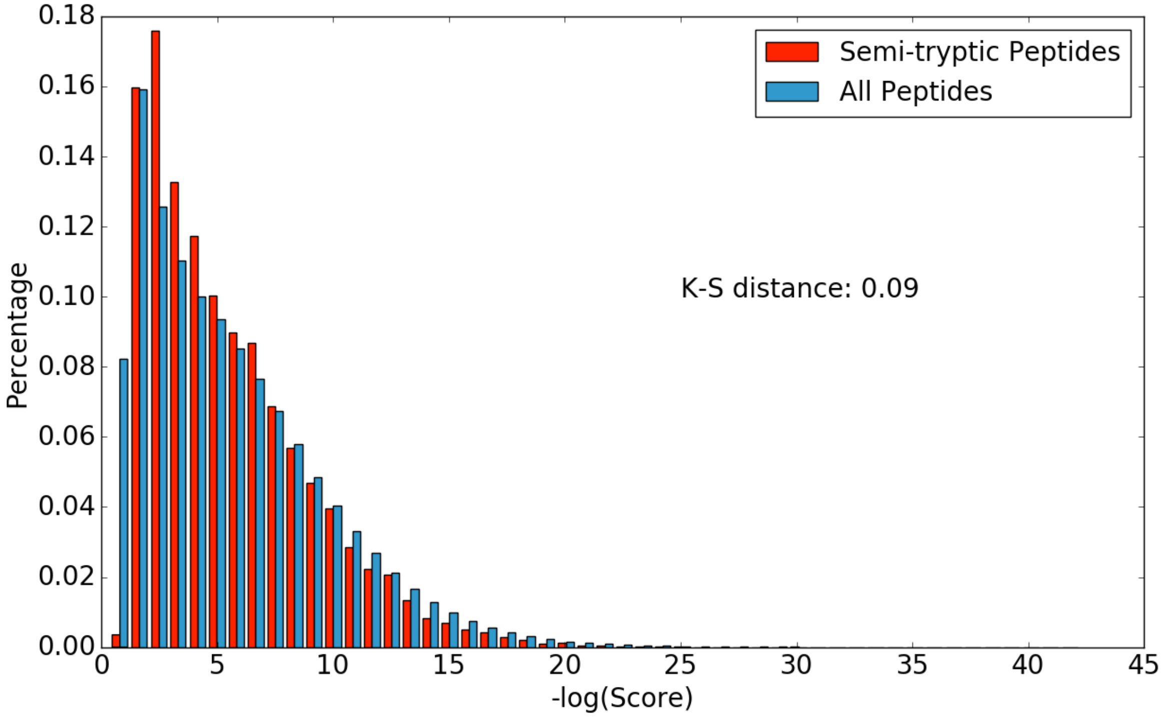

**Supplementary Fig. 9.** The score distributions from the 9,559 semi-tryptic peptides and the scores of all 548,371 peptides identified in the Kim data.

**a**

Estimated Accuracy of the consistently identified PSMs (%)

| Row. ∩ Col. | Open-pFind | MSFragger | PEAKS | MODa | pFind | MS-GF+ | Comet | Byonic |
|---|---|---|---|---|---|---|---|---|
| Open-pFind | 99.2 | 99.9 | 99.2 | 100.0 | 99.4 | 99.4 | 99.8 | 99.5 |
| MSFragger | 99.9 | 98.2 | 99.9 | 98.9 | 99.9 | 99.2 | 99.0 | 99.9 |
| PEAKS | 99.2 | 99.9 | 98.4 | 99.9 | 99.4 | 99.3 | 99.7 | 99.4 |
| MODa | 100.0 | 98.9 | 99.9 | 98.6 | 100.0 | 99.2 | 99.1 | 99.9 |
| pFind | 99.4 | 99.9 | 99.4 | 100.0 | 99.2 | 99.4 | 99.9 | 99.5 |
| MS-GF+ | 99.4 | 99.2 | 99.3 | 99.2 | 99.4 | 95.7 | 97.0 | 99.5 |
| Comet | 99.8 | 99.0 | 99.7 | 99.1 | 99.9 | 97.0 | 95.9 | 99.9 |
| Byonic | 99.5 | 99.9 | 99.4 | 99.9 | 99.5 | 99.5 | 99.9 | 97.5 |

Estimated Accuracy of the separately identified PSMs (%)

| Row. – Col. | Open-pFind | MSFragger | PEAKS | MODa | pFind | MS-GF+ | Comet | Byonic |
|---|---|---|---|---|---|---|---|---|
| Open-pFind | | 98.6 | 99.1 | 98.6 | 99.0 | 98.9 | 98.5 | 98.9 |
| MSFragger | 85.3 | | 92.7 | 96.3 | 93.7 | 95.2 | 95.4 | 93.3 |
| PEAKS | 90.9 | 96.3 | | 96.3 | 94.4 | 95.7 | 93.9 | 95.4 |
| MODa | 81.5 | 96.9 | 89.9 | | 93.5 | 95.9 | 96.2 | 92.9 |
| pFind | 95.8 | 98.0 | 97.0 | 98.0 | | 98.1 | 94.9 | 98.0 |
| MS-GF+ | 63.3 | 88.5 | 68.2 | 89.4 | 73.0 | | 77.4 | 81.4 |
| Comet | 66.1 | 89.8 | 72.1 | 90.7 | 75.2 | 87.7 | | 82.4 |
| Byonic | 82.5 | 92.0 | 81.0 | 92.6 | 85.9 | 88.4 | 84.5 | |

**b**

Estimated Accuracy of the consistently identified PSMs (%)

| Row. ∩ Col. | Open-pFind | MSFragger | PEAKS | MODa | pFind | MS-GF+ | Comet | Byonic |
|---|---|---|---|---|---|---|---|---|
| Open-pFind | 98.9 | 99.7 | 99.6 | 99.9 | 99.8 | 99.8 | 99.7 | 99.9 |
| MSFragger | 99.7 | 86.6 | 99.0 | 97.6 | 99.9 | 99.1 | 99.0 | 99.9 |
| PEAKS | 99.6 | 99.0 | 93.5 | 98.5 | 99.4 | 99.2 | 99.6 | 99.4 |
| MODa | 99.9 | 97.6 | 98.5 | 90.0 | 100.0 | 99.5 | 99.3 | 100.0 |
| pFind | 99.8 | 99.9 | 99.4 | 100.0 | 99.2 | 99.4 | 99.9 | 99.5 |
| MS-GF+ | 99.8 | 99.1 | 99.2 | 99.5 | 99.4 | 95.7 | 97.0 | 99.5 |
| Comet | 99.7 | 99.0 | 99.6 | 99.3 | 99.9 | 97.0 | 95.9 | 99.9 |
| Byonic | 99.9 | 99.9 | 99.4 | 100.0 | 99.5 | 99.5 | 99.9 | 97.5 |

Estimated Accuracy of the separately identified PSMs (%)

| Row. – Col. | Open-pFind | MSFragger | PEAKS | MODa | pFind | MS-GF+ | Comet | Byonic |
|---|---|---|---|---|---|---|---|---|
| Open-pFind | | 98.2 | 98.0 | 98.2 | 98.2 | 98.2 | 98.2 | 98.3 |
| MSFragger | 45.2 | | 65.3 | 72.1 | 71.3 | 69.7 | 68.5 | 73.6 |
| PEAKS | 63.9 | 87.1 | | 87.8 | 82.1 | 83.0 | 81.6 | 84.5 |
| MODa | 23.2 | 72.6 | 48.5 | | 67.7 | 60.5 | 60.1 | 70.1 |
| pFind | 87.9 | 97.3 | 96.9 | 97.8 | | 98.1 | 94.9 | 98.0 |
| MS-GF+ | 76.1 | 89.2 | 83.4 | 91.9 | 73.0 | | 77.4 | 81.4 |
| Comet | 83.4 | 91.8 | 84.5 | 93.4 | 75.2 | 87.7 | | 82.4 |
| Byonic | 80.6 | 91.5 | 80.5 | 93.0 | 85.9 | 88.4 | 84.5 | |

**Supplementary Fig. 10.** The estimated accuracy of consistently and separately identified PSMs from the comparison of every two search engines using the Dong-Ecoli-QE dataset. $^{15}N$- and $^{13}C$-labeled peptides are used for estimation, and the final accuracy is calculated from the average of the two estimates for the same resulting PSMs. Each decimal denotes the estimated accuracy of the consistently or separately identified PSMs. a) Only the PSMs with common modification types (the four that are specified in the restricted search engines) are considered in the statistical analysis. b) All PSMs are considered in the statistical analysis.
