## Supplementary Materials for "Open-pFind enables precise, comprehensive and rapid peptide identification in shotgun proteomics"

### Supplementary Note 1

**An overview of the Open-pFind results from the Dong-Ecoli-QE dataset.** As shown in Fig. S1a, the precursor mass deviations of most identified peptides were distributed within ±3 ppm. Thus, there were scarcely any incorrect peptides with abnormal mass deviations based on high resolution MS. The numbers of target and decoy PSMs with lower scores were essentially the same, which was consistent with the hypothesis that the ratio between the target and decoy random matches must be approximately 1:1 (Fig. S1b). There were ~90% fully specific peptides, while the proportion of non-specific peptides was less than 1% (Fig. S1c), and the missed cleavage sites of over 99% of peptides were less than two (Fig. S1d). In addition, only six modifications were abundantly discovered in over 1% of peptides, all of which were commonly specified in routine MS/MS data analyses (Fig. S1e). Finally, ~13.5% spectra were identified with more than one peptide, while most yielded two resulting peptides (Fig. S1f).

A mixed spectrum of three co-eluting peptides is shown as an example in Fig. S2. Three isotopic clusters are clearly distinguished from each other, while the observed fragmentation sites were near-complete for all three peptides. Notably, most of the fragment ions were uniquely matched to one of the three peptides. Furthermore, the fragmentation patterns of the three peptides were predicted using pDeep[1]. The Pearson's correlation coefficients (PCCs) between the actual PSM and the predicted PSM were as high as 0.97, 0.99 and 0.87 for the three peptides (Fig. S3), suggesting the high accuracy of Open-pFind for the analysis of mixed spectra.

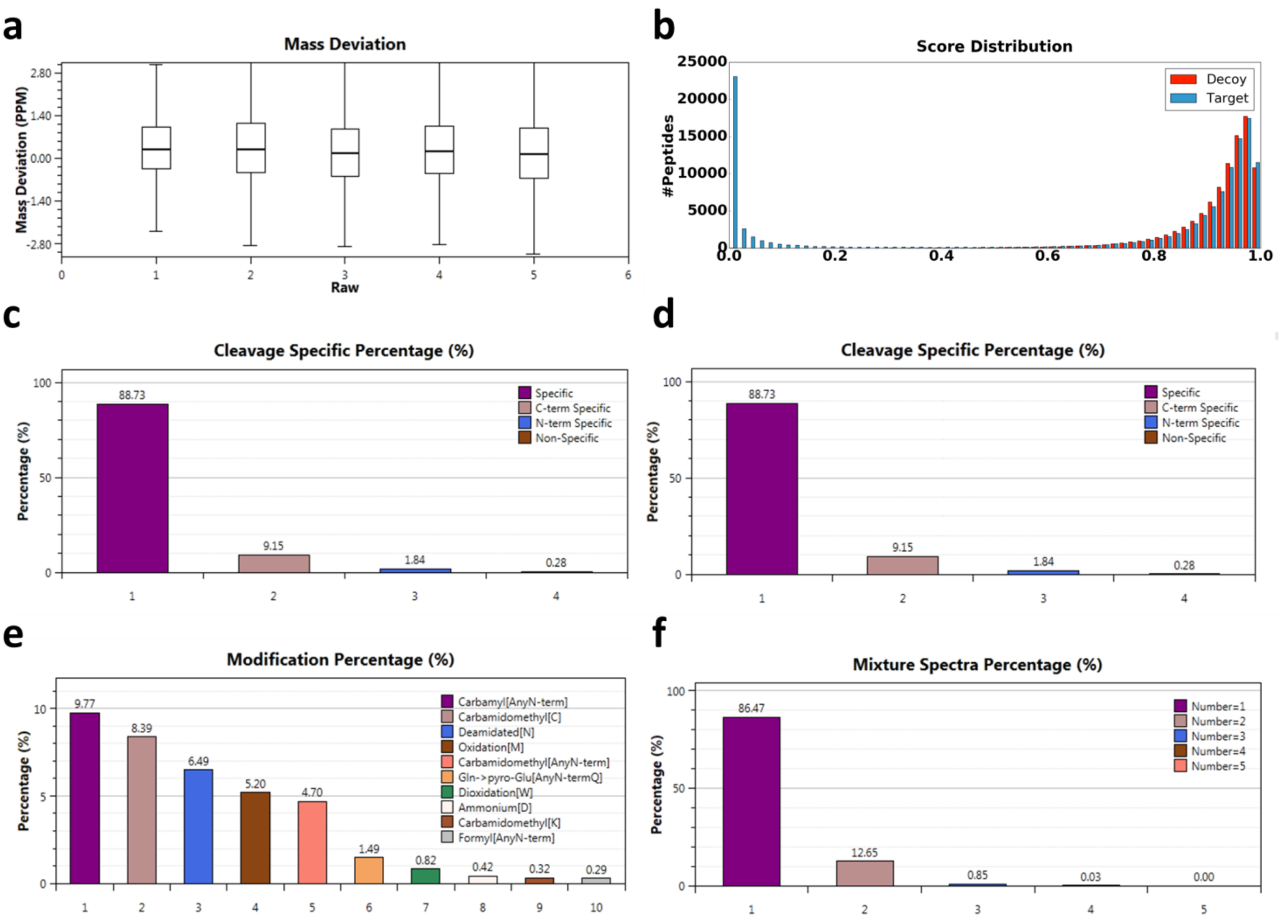


**Fig. S1.** An overview of the results reported by Open-pFind for the Dong-Ecoli-QE dataset. This figure is a screenshot of the in-house software tool pBuild, which was designed to graphically represent pFind Studio results. **a)** The distributions of mass deviations (in ppm) of the identified PSMs in the five raw files. **b)** Score distributions of the target (blue) and decoy (red) PSMs. **c)** Distribution of the enzymatic specificity of identified peptides. **d)** Percentages of the highly abundant modifications in all identified peptides. **e)** Distribution of the number of missed cleavage sites. **f)** Distribution of the number of peptides identified from one tandem mass spectrum. For example, there are 12.65% spectra, each of which yielded two peptides.

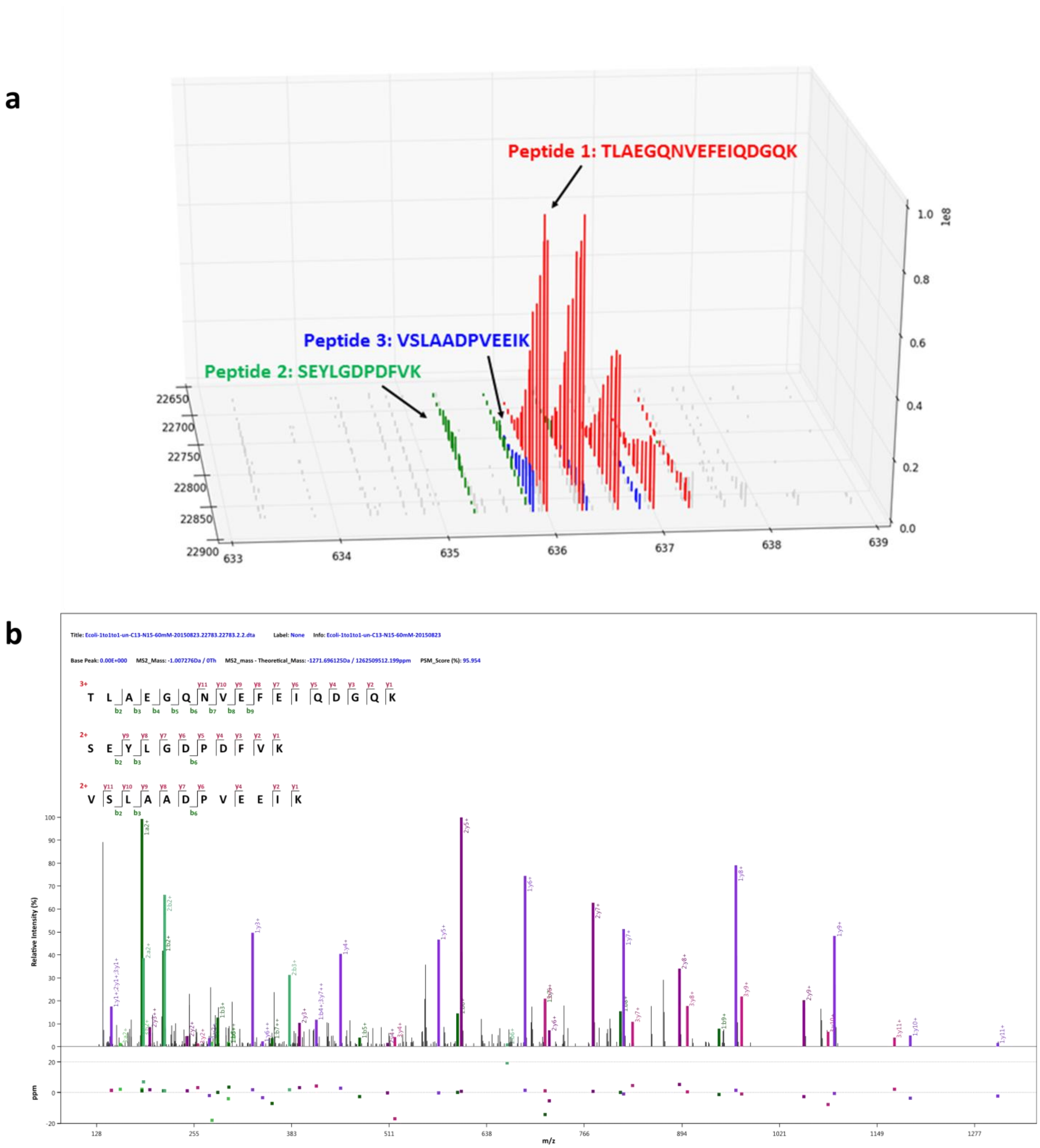


**Fig. S2.** A spectrum entitled “Ecoli-1to1to1-un-C13-N15-60mM-20150823.22783.22783.dta”, in which three co-eluting peptides were identified. **a)** MS1 information for the three peptides in the isolation window around this tandem mass spectrum. **b)** The match between the spectrum and the three resulting peptides, taken from a screenshot of pBuild.

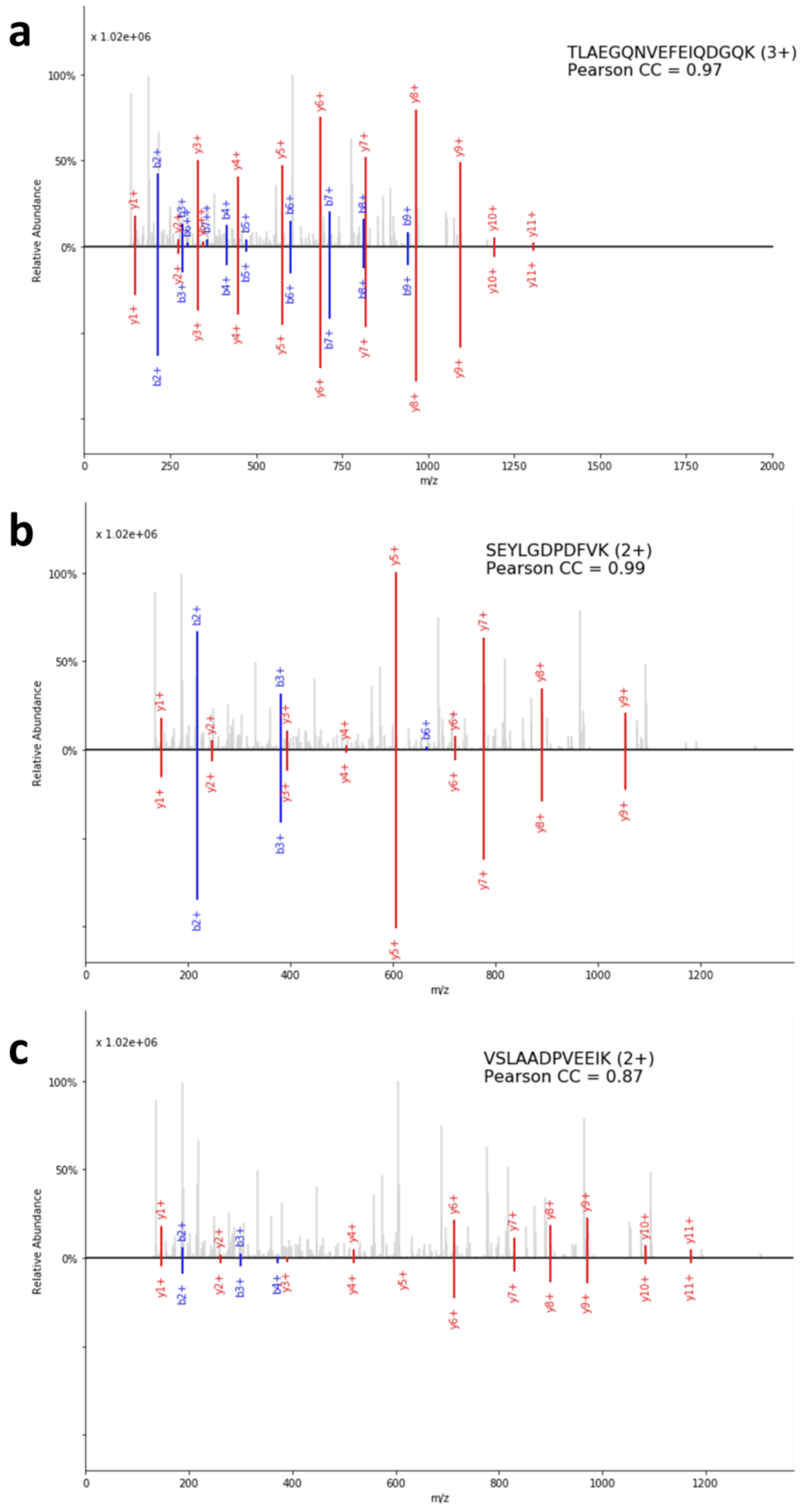


**Fig. S3.** Similarity between the actual and predicted fragment ions for each of the three co-eluting peptides identified from Ecoli-1to1to1-un-C13-N15-60mM-20150823.22783.22783.2.dta and measured with PCC: **a)** TLAEGQNVEFEIQDGQK, **b)** SEYLGDPDFVK and **c)** VSLAADPVEEIK.

1. Zhou, X.X. et al. pDeep: Predicting MS/MS Spectra of Peptides with Deep Learning. *Anal Chem* (2017).

### Supplementary Note 2

**Comparison of Open-pFind, MaxQuant and SEQUEST-HT showed the advantages of Open-pFind, both in terms of sensitivity and accuracy.** Open-pFind was compared with MaxQuant and SEQUEST-HT in terms of consistency and the proportion of NaN-ratio PSMs. Open-pFind covered over 90% of the results of MaxQuant and SEQUEST-HT and provided 54.3% of the identifications separately (Fig. S4a). Furthermore, for the intersecting results from the three engines, the NaN-ratio PSMs constituted a smaller fraction than those separately identified by each engine (Fig. S4b). The proportion of NaN-ratio PSMs in the separate results from Open-pFind was the smallest, indicating that the PSMs identified by Open-pFind were more accurate than the other two search engines. Additionally, we compared the running times of these three engines using the same configuration (Supplementary Table 3), and Open-pFind was 3.4 and 15.7 times faster than SEQUEST-HT and MaxQuant, respectively (Fig. S5).

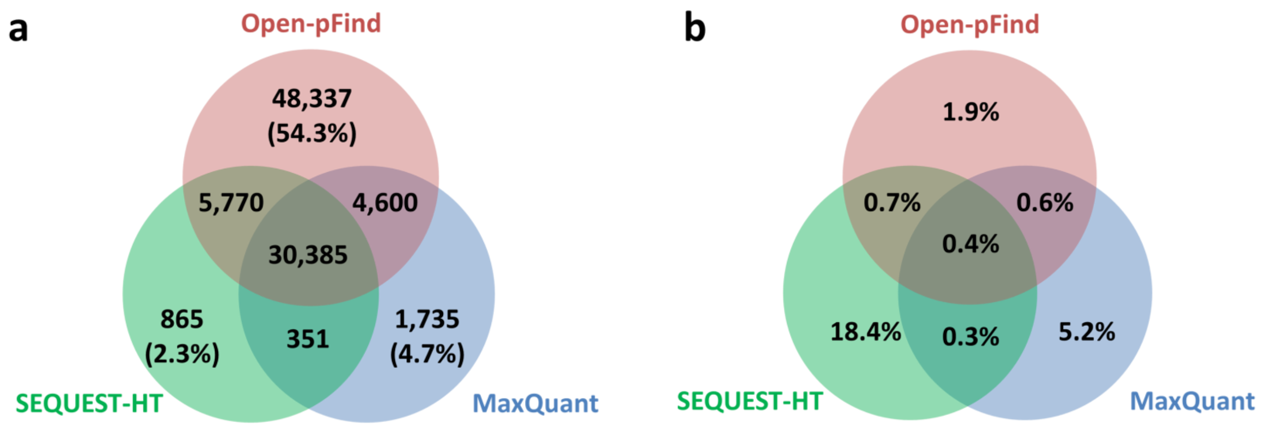


**Fig. S4.** Comparison of the results from Open-pFind, SEQUEST-HT and MaxQuant. **a)** The consistency of the identified PSMs. Each of the numbers in parentheses denotes the percentage of uniquely identified PSMs in the total results. **b)** The percentages of NaN-ratio PSMs in different parts of the results.

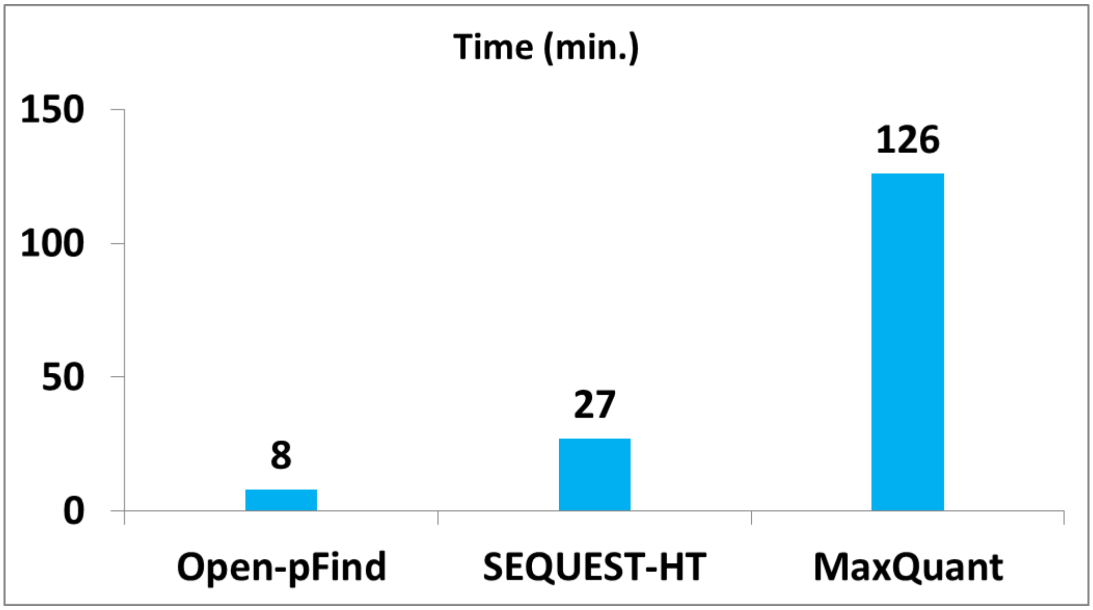


**Fig. S5.** Running times for Open-pFind, SEQUEST-HT and MaxQuant.

### Supplementary Note 3

**Analysis based on the entrapment strategy showed the robustness of the design of Open-pFind.** To analyze four published datasets, two types of entrapment databases were downloaded from the UniProt database and then used in this study: a) a small database of the reviewed proteins of *Arabidopsis thaliana* (8.7 MB, 15,423 protein sequences) and b) a large database of the reviewed proteins of all organisms (261.8 MB, 555,100 protein sequences). The entrapment databases were appended to the original database files, respectively. The other database search parameters were the same as those shown in Supplementary Table 3.

Intuitively, when the entrapment database is considered in the database search, the identification rate should decrease because more random peptide candidates are involved in the search space, but few of them are the answers to any spectra. Generally, the decrease was more remarkable when the larger entrapment database was considered (Fig. S6). The Open-pFind identification rate was more stable in both situations than that of pFind. For example, the average decrease in the identification rates of Open-pFind and pFind was 1.6 and 4.2, respectively (Fig. S6b), because Open-pFind adopted a two-step workflow, and the proteins to be retrieved in the restricted search were automatically learned in the previous open search step; thus, most random peptide candidates that potentially interfere with the correct candidates were eliminated at this time. Furthermore, only less than 5% of PSMs from Open-pFind that matched with the entrapment sequences were revived by Open-pFind (Fig. S7); however, the corresponding pFind percentages varied from 20% to 60%. This phenomenon was similar to that observed with the NaN-ratio analysis of the Dong-Ecoli-QE dataset, which again proved that Open-pFind reported more accurate peptides that matched the authentic protein sequences rather than the entrapment sequences, although the same FDR threshold was controlled.

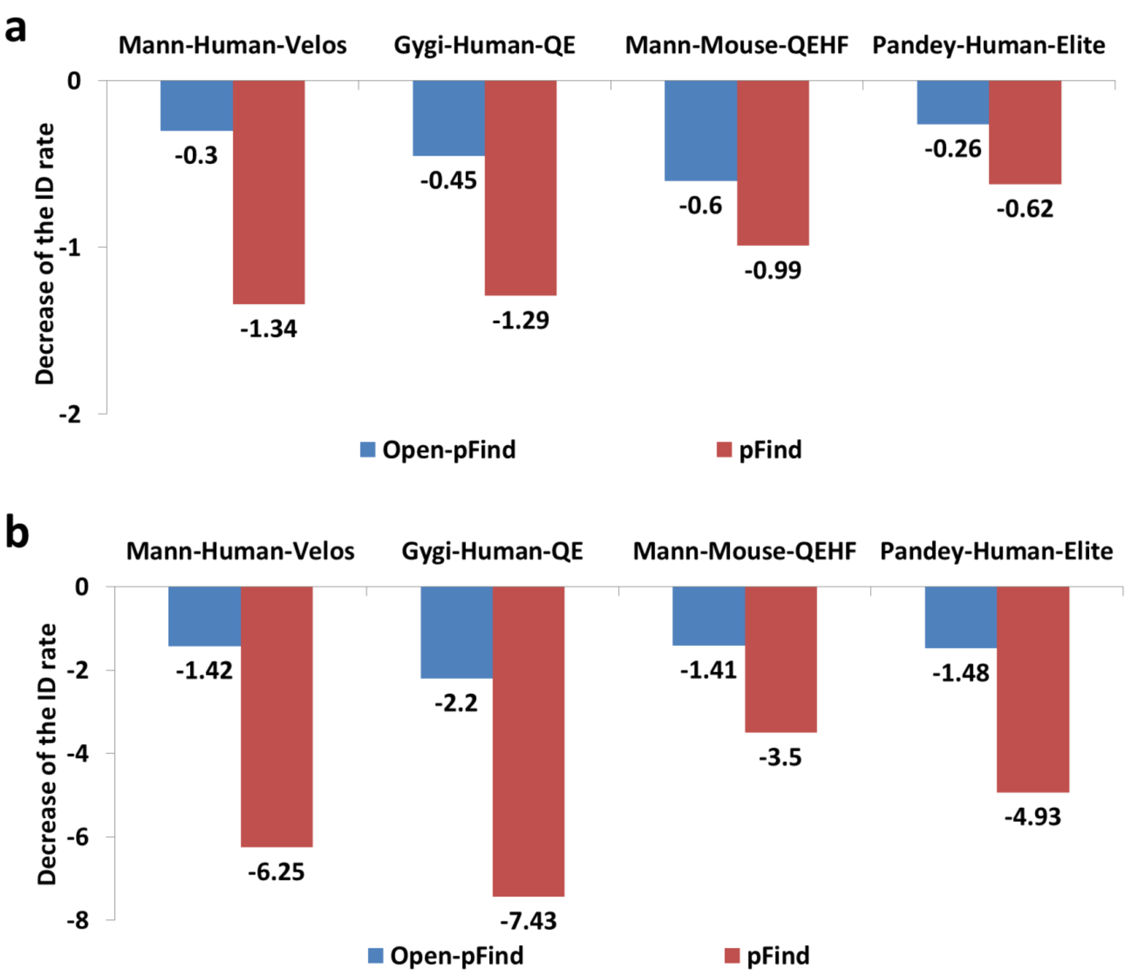


**Fig. S6.** Decreased identification rates caused by the entrapment strategy for the four datasets. **a)** Proteins from *Arabidopsis thaliana* were considered the entrapment database. **b)** Proteins from all organisms recorded in UniProt were considered the entrapment database.

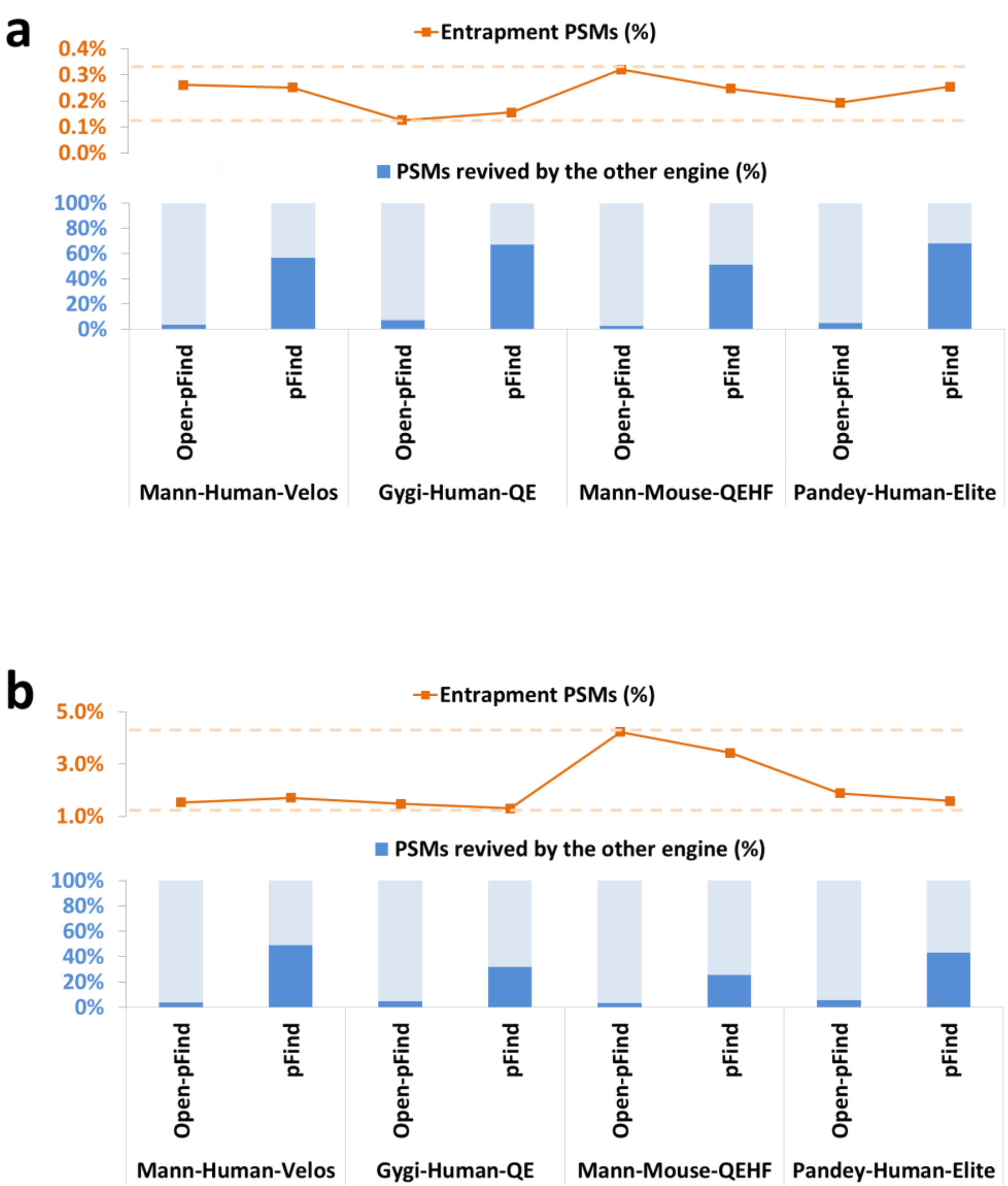


**Fig. S7.** Open-pFind revived more spectra than pFind. The orange curves denote the proportion of PSMs from the entrapment database. **a)** Proteins from *Arabidopsis thaliana* were considered the entrapment database. b) Proteins from all organisms recorded in UniProt were considered the entrapment database.

We also used the entrapment strategy to evaluate the precision of search engines with the Dong-Ecoli-QE dataset (the reviewed human database downloaded from UniProt was used as the entrapment database), and the performance of Open-pFind was similar to that of the four large-scale datasets. When searching against the target and entrapment databases, Open-pFind reported the highest numbers of PSMs with the smallest proportions of those matched with the entrapment proteins (Fig. S8a). Only less than 10% of PSMs from Open-pFind that matched the entrapment proteins were revived by the other engines, while 22–56% of the PSMs from other engines were revived by Open-pFind (Fig. S8b).

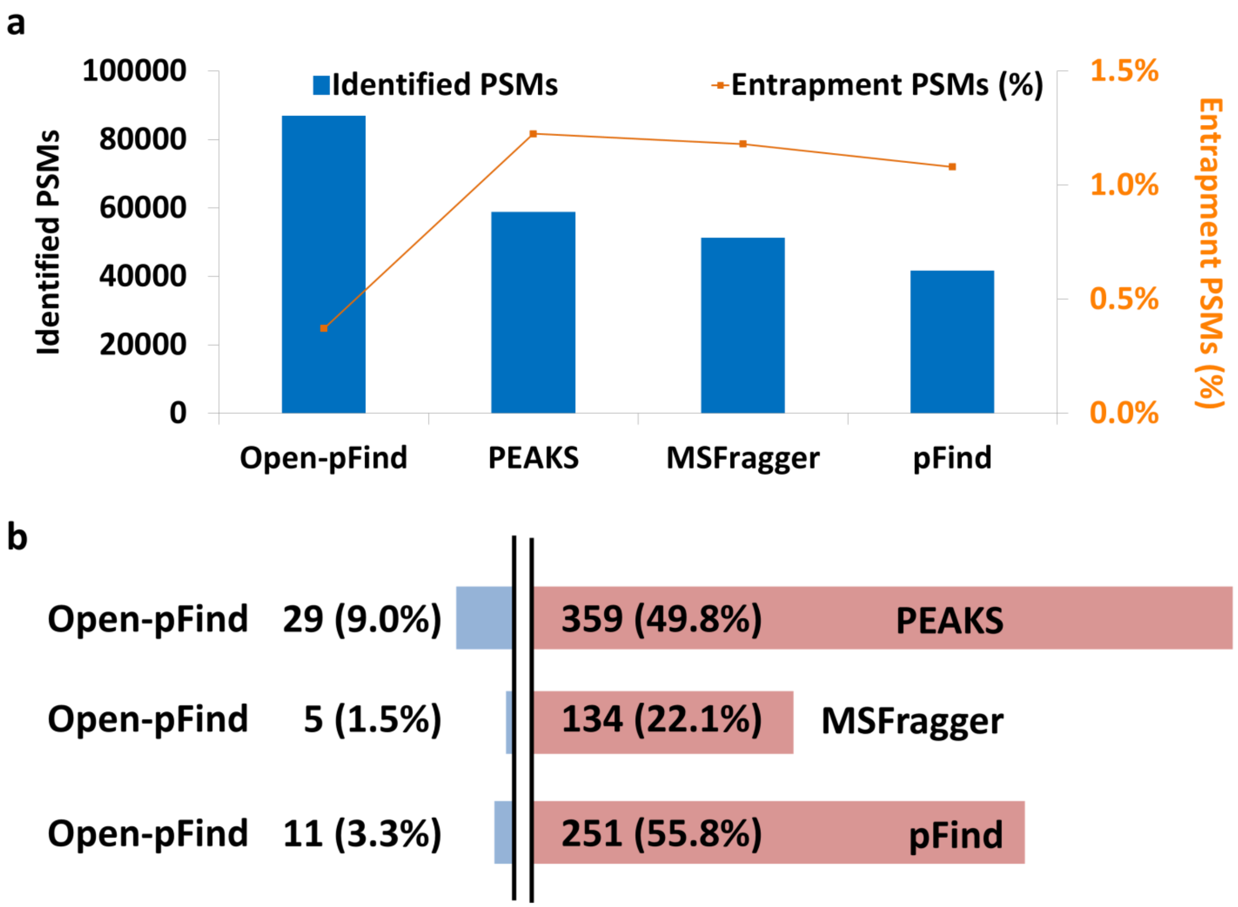


**Fig. S8.** Entrapment analysis of the Dong-Ecoli-QE dataset. a) The number of identified PSMs (the blue bars) and the percentage of PSMs from the entrapment database (the orange curve). b) The number and proportion of PSMs identified with entrapment peptides from one engine and revived by the other engine.

### Supplementary Note 4

**Using NaN ratios to estimate the error rates of different engines independent of the target-decoy strategy.** First, we investigated the relationship between decoy PSMs and NaN-ratio PSMs based on the Open-pFind results obtained for the Dong-Ecoli-QE dataset. Fig. S9 shows the increase in the number of decoys and NaN-ratio PSMs along with the numbers of target PSMs (all PSMs were sorted in descending order of their $q$-values). The trends of the three curves were quite consistent, and the tails (where nearly all PSMs were incorrect) showed that the proportions of both decoy and NaN-ratio PSMs were stable.

Therefore, the percentage of NaN-ratio PSMs is useful for estimating the error rates of the results of metabolically-labeled datasets, which is similar to but independent of the traditional target-decoy strategy. Given $M$ as the number of total PSMs and $N$ as the number of NaN-ratio PSMs, we get the equation

$$M * e * r_1 + M * (1 - e) * r_2 = N, \quad 1)$$

where $e$ denotes the error rate to be estimated, $r_1$ denotes the percentage of NaN-ratio PSMs in *incorrect* matches (*e.g.*, target PSMs distributed at the tail of the curves in Fig. S9) and $r_2$ denotes the percentage of NaN-ratio PSMs in *correct* matches. $r_1$ is simply calculated using the linear least-squares method, and $r_2$ is estimated based on the intersection of the results of different engines because a PSM is more likely to be correct if it is consistently reported by multiple search engines, resulting in a lower probability of being a NaN-ratio PSM (Fig. S10). In this study, the intersecting results of all eight search engines were used to estimate the value of $r_2$ (Fig. S11). Finally, the error rate $e$ is estimated using the following formula:

$$e = \frac{N - M * r_2}{M * (r_1 - r_2)}, \quad 2)$$

and the accuracy of the given result set is equal to $1 - e$.

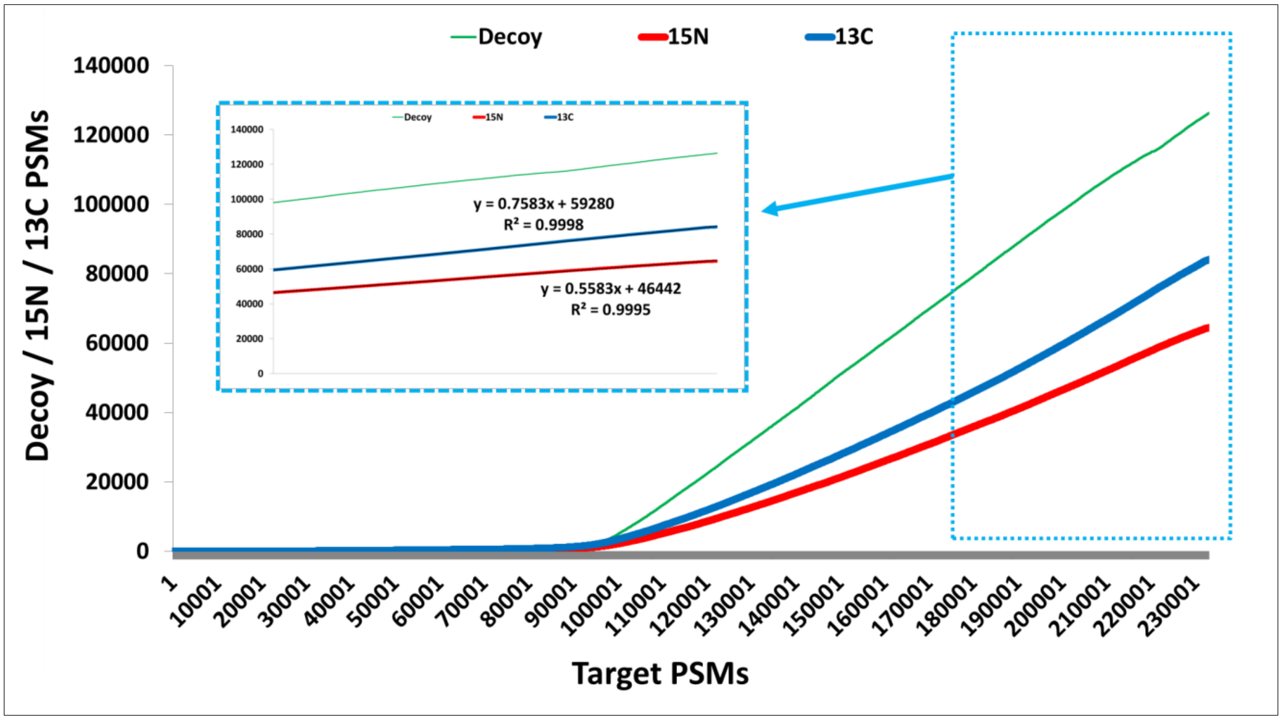


**Fig. S9.** The number of PSMs from the decoy database (green) or with NaN ratios of $^{15}N/^{14}N$ (red) or $^{13}C/^{12}C$ (blue) corresponding to the number of target PSMs in the Dong-Ecoli-QE dataset. The subplot shows where the linear property is used to estimate the proportion of NaN-ratio PSMs.

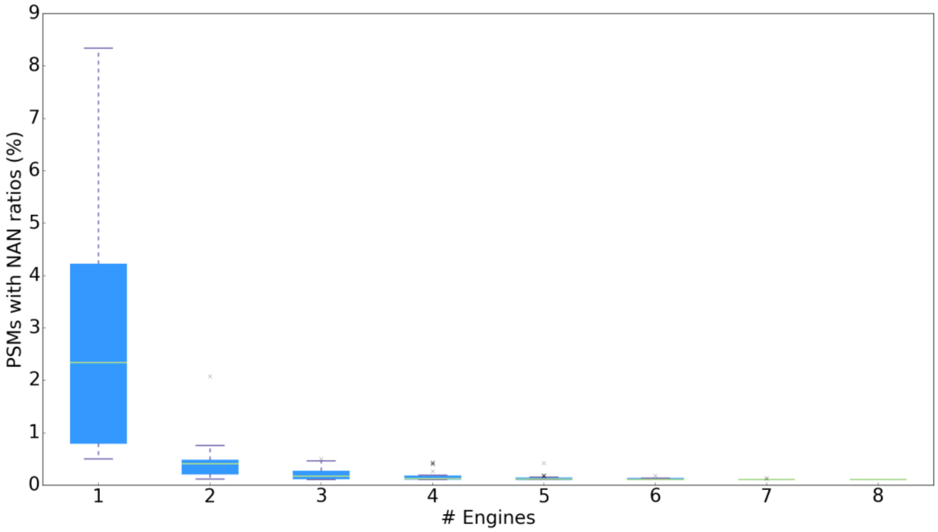


**Fig. S10.** The proportions of NaN-ratio PSMs distributed in all of the possible intersections of the eight result sets from Open-pFind, PEAKS, MODa, MSFragger, MS-GF+, Byonic, Comet and pFind. For example, the number of intersections from any three result sets is $\binom{8}{3} = 56$.
